# AF2χ: Predicting protein side-chain rotamer distributions with AlphaFold2

**DOI:** 10.1101/2025.04.16.649219

**Authors:** Matteo Cagiada, F. Emil Thomasen, Sergey Ovchinnikov, Charlotte M. Deane, Kresten Lindorff-Larsen

## Abstract

The flexibility of protein side chains is an essential contributor of conformational entropy and affects processes such as folding, stability and molecular interactions. Structure determination experiments and prediction tools such as AlphaFold generally fail to capture or represent the conformational heterogeneity of proteins in solution. Experiments can be used to study side-chain flexibility, but cannot be applied at scale, and most prediction methods focus on reconstructing the minimum free energy state rather than an ensemble representing side-chain configurations. Here, we use AlphaFold2 and its internal side-chain representations to develop AF2χ that predicts side-chain χ-angle distributions and generates structural ensembles. We extensively benchmark AF2χ predictions using experimental NMR ^3^*J*-couplings and *s*^2^ order parameters, as well as dihedral angle distributions derived from collections of experimental structures, demonstrating the accuracy of AF2χ in generating accurate side-chain ensembles. We also compare the accuracy of AF2χ with molecular dynamics simulations and recent machine learning models aimed to generate conformational ensembles and show that AF2χ provides state-of-the-art accuracy orders of magnitude faster than molecular simulations. With its speed and accuracy, AF2χ offers a strong complementary option to simulations and rotamer library approaches, making it particularly valuable for applications such as protein design, ligand docking and interpretation of biophysical experiments.

## Introduction

The flexibility of protein side chains is important for conformational entropy and plays a crucial role in processes such as protein folding (***Ohgushi and Wada, 1983***; ***Furukawa et al., 1996***), stability (***Frauenfelder et al., 1991***; ***Doig and Sternberg, 1995***; ***Mittermaier and Kay, 2004***) and molecular interactions (***Rasmussen et al., 1992***; ***Eisenmesser et al., 2002***; ***Zavodszky and Kuhn, 2005***; ***Frederick et al., 2007***; ***Wankowicz and Fraser, 2024***). The flexibility of side chains can vary substantially across a protein (***Mittermaier et al., 1999***; ***Marlow et al., 2010***), with some residues being rigid where side-chain dihedral angles (χ-angles) are highly constrained by their local environment to populate mostly a single rotamer state, whereas others show substantial heterogeneity (***Schrauber et al., 1993***). This heterogeneity depends on the specific environment, and is influenced by factors such as local backbone structure and the number of contacts which the side chain makes (***Best et al., 2004***; ***Ming and Brüschweiler, 2004***).

Conventionally used for the determination of static protein structures (Fig. 1a,b), experimental techniques such as X-ray crystallography (***Best et al., 2006***; ***Fraser et al., 2011***; ***Fenwick et al., 2014***), cryo-electron microscopy (cryo-EM) (***Riley et al., 2021***; ***Hoff et al., 2024***), and nuclear magnetic resonance (NMR) spectroscopy (***Kay, 1998***; ***Mittermaier and Kay, 2001***) can also provide information about the conformational heterogeneity of side chains. For example, room-temperature X-ray crystallography can provide information on side-chain structural heterogeneity directly in the electron density (***Fraser et al., 2011***; ***Fenwick et al., 2014***), NMR relaxation experiments can provide information on the conformational dynamics of side-chains, and NMR scalar (^3^*J*) and residual-dipolar couplings can provide ensemble-averaged information about χ-angles (***Palmer III, 2004***). Experiments that report on side-chain flexibility can also be used in combination with computational protocols, such as Dynamic Ensemble Refinement (DER) (***Lindorff-Larsen et al., 2005a***) where molecular dynamics (MD) simulations are refined against data from solution NMR, leading to accurate representations of side-chain conformational heterogeneity in solution (Fig. 1c,d).

**Figure 1.**
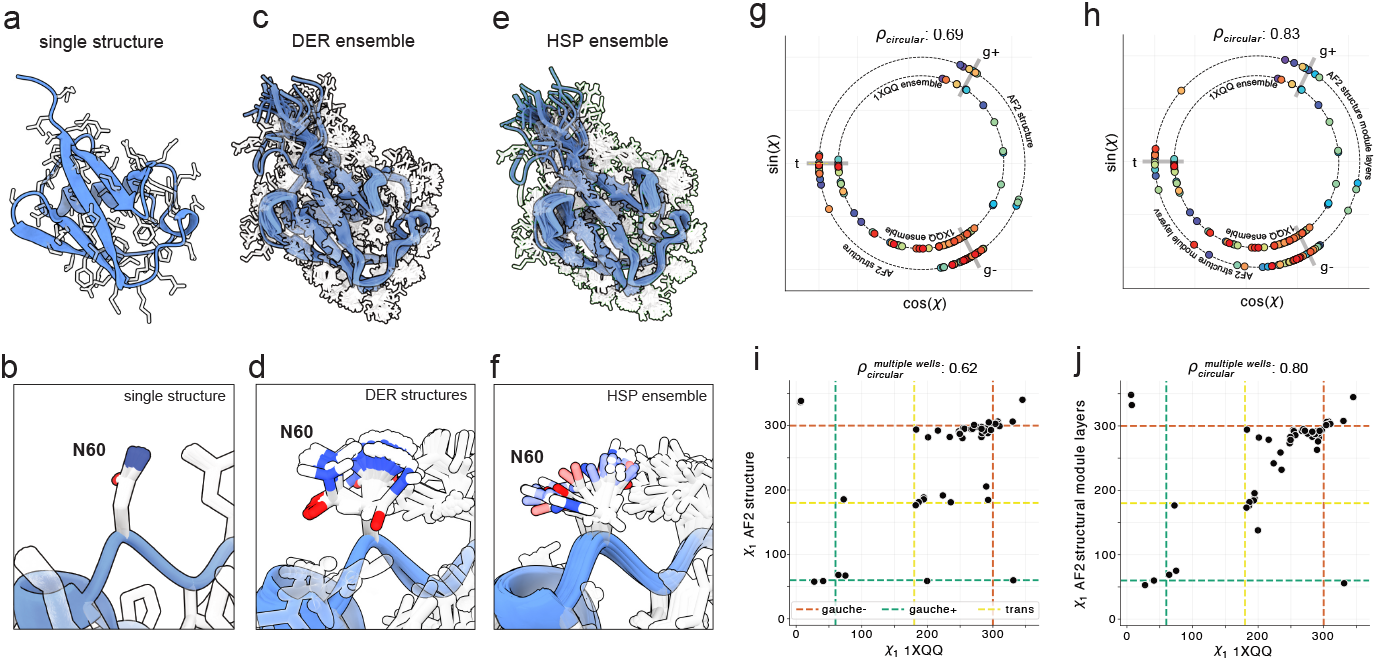
Capturing side-chain conformational heterogeneity using structural ensembles. **(a-f)** Illustration of the side-chain flexibility in different structural models of UBQ. The structures show secondary structure elements as navy blue cartoon and side-chains as transparent white atoms. The bottom panels show the same residue in the different structural models. **(a**,**b)** A single structure obtained using X-ray diffraction experiments (PDB ID: 1UBI). **(c**,**d)** A DER ensemble obtained from MD simulations restrained by NMR data (PDB: 1XQQ). **(e**,**f)** An HSP ensemble obtained by overlaying 88 PDB structures. **(g**,**h)** Average χ_1_ angles from the DER ensemble (PDB: 1XQQ) (inner circle) and the χ_1_-angles from the AF2 (g) outer or (h) inner layer (outer circle). **(i**,**j)** Correlation between the average χ_1_ angles from the DER ensemble and the χ_1_-angles from the AF2 (i) inner and (j) outer layers for the subset of residues that sample multiple rotamer states. In (g) and (h) each residue has a unique colour, paired between the DER ensemble value (inner circle) and the AF2 value (outer circle). The three staggered rotamer states for χ_1_ are shown with letters and a line in (g) and (h) and as dotted lines in panels (i) and (j).

Structure-based rotamer libraries have become a widely used tool to predict side-chain configurations. Rotamer libraries catalogue the distribution of side-chain dihedral angles observed in high-quality protein structures into discrete distributions. Such rotamer libraries are used in several computational tasks, like modelling the side-chain structure and dynamics (***Canutescu et al., 2003***; ***Bhowmick and Head-Gordon, 2015***), protein-protein docking (***Wang et al., 2005***), homology modeling, and protein design (***Havranek and Baker, 2009***). Over the years, rotamer libraries have been improved by implementing increasingly stringent selection criteria to ensure accuracy and diversity of datasets, as well as adding backbone context using different clustering methods (***Dunbrack and Karplus, 1993***; ***Lovell et al., 2000***; ***Hintze et al., 2016a***). This has improved side-chain representations in specific protein environments, such as different secondary structure elements. However, rotamer libraries were not designed to represent side-chain heterogeneity. Instead, they help categorise the types of structures that are commonly seen, but not necessarily the distribution of conformations present at a given site in solution.

Information on side-chain flexibility can be obtained from the variation across static protein structures with high-sequence similarity, known as High Sequence Similarity Protein Data Bank (HSP) ensembles (Fig. 1e,f) (***Best et al., 2006***). The idea behind HSP ensembles is that structural perturbations caused by differences in e.g. crystal environment, temperature, ligand binding, or mutations help expose the equilibrium structural fluctuations in solution. MD simulations have also been used extensively to study heterogeneity of side chain conformations (***Das and Baker, 2008***; ***Lindorff-Larsen et al., 2010***; ***Bowman and Geissler, 2014***; ***Petrović et al., 2018***), but due to their computational cost, MD simulations cannot easily be scaled up to large numbers of different systems.

Recent advances in deep learning have given us the ability to predict protein structure using tools such as AlphaFold2 (AF2) (***Jumper et al., 2021***), and to model side-chain rotamers given an input backbone using tools such as DLpacker (***Misiura et al., 2022***) or H-packer (***Visani et al., 2023***). These tools are, however, not designed for studying side-chain heterogeneity, as they typically aim to recover the major dihedral states given an input backbone, using static X-ray crystal structures as their benchmark for accuracy.

Here we present AF2χ, a prediction model based on AF2 that generates a collection of side-chain dihedral distributions for each residue, and then a set of conformations, i.e. a structural ensemble, conditioned on the dihedral distributions. We extensively validate AF2χ outputs against experimental data such as HSP ensembles, NMR *s*^2^ order parameters, and ^3^*J*-couplings, as well as MD simulations. We also compare the accuracy of AF2χ to other methods for predicting conformational dynamics. With its speed and accuracy, AF2χ offers a powerful complement to MD simulations and rotamer libraries for protein design, ligand docking, and biophysical analyses.

## Results and Discussion

Our goal with AF2χ was to develop a method that could rapidly and accurately model the conformational heterogeneity of the side chains in a protein structure in solution, and to represent this with an ensemble of conformations. Our work thus complements recent work that has aimed at predicting protein dynamics at the level of the polypeptide backbone. In addition to developing AF2χ we also aimed to collect a broad set of experimental and computational data that we could use to benchmark our ability to predict solution-state dynamics.

### HSP ensembles

HSP ensembles are generated by overlaying experimental protein structures with high sequence-similarity. They have been shown to accurately reproduce side-chain dynamics in solution (***Best et al., 2006***). Building on this previous work, we first tested whether HSP ensembles can be used to accurately predict side-chain dihedral angle distributions. We tested the accuracy of HSP ensembles in reproducing NMR scalar ^3^*J*-couplings that report on χ_1_ dihedral angles. ^3^*J*-couplings are ensemble-averaged, and are therefore sensitive to the distribution of rotamer states in solution. We identified three systems for which ^3^*J*-couplings probing χ_1_ angles have been reported, and for which sufficient experimental structures have been determined to create large HSP ensembles: ubiquitin (UBQ), hen egg white lysozyme (HEWL) and bovine pancreatic trypsin inhibitor (BPTI). We constructed HSP ensembles of these proteins using structures from the PDB (***Berman et al., 2000***) and calculated ^3^*J*-couplings corresponding to the experimental data using Karplus equations. We found a good correlation with experimental ^3^*J*-couplings (Fig. S1). For UBQ, we also calculated ^3^*J*-couplings from a previously reported DER ensemble, which was determined based on experimental *s*^2^ order parameters and nuclear Overhauser effect data and validated with residual dipolar couplings and ^3^*J*-couplings (***Lindorff-Larsen et al., 2005a***). We found that the correlation to ^3^*J*-couplings given by our HSP ensemble matches the accuracy of the DER ensemble (Fig. S1a). As previously shown based on the correlation with NMR order parameters (***Best et al., 2006***), the ability of HSP ensembles to reproduce native state dynamics decreases rapidly when fewer structures are available. To further test this, we used a bootstrap sampling procedure to generate HSP ensembles of UBQ, HEWL, and BPTI with a varying number of structures. We compared the resulting ensembles with ^3^*J*-couplings and found that at least 20 structures are required to achieve a good correlation (Fig. S1b,d,f), consistent with what was previously found when comparing with NMR *s*^2^ order parameters (***Best et al., 2006***). This means that HSP ensembles are not a generalizable approach to characterizing the structural heterogeneity of protein side chains, as most proteins do not have 20 or more experimental structures.

### Information on rotamer distributions from the AF2 structure module

We next tested whether deep learning structure prediction tools such as AF2 (***Jumper et al., 2021***) could be used to predict protein side-chain flexibility as an alternative to HSP ensembles. As AF2 explicitly predicts χ-angle values for side chains, we hypothesised that it may contain information about side-chain structural heterogeneity. Specifically, we asked the question whether we could extract such information from the side-chain unit of the structure module, where the final χ values are generated. Using UBQ as a test system, we extracted χ_1_ and χ_2_ predictions from the eight different layers of the multi-rigid side-chain unit. We observed that the predicted χ-angle changed very little throughout the first seven layers, especially for χ_1_, whereas the final layer produced more substantial changes in the predicted χ-angle for certain residues (Fig. S2a,b). Additionally, we found that while the inner layers in some cases predict χ-angles that are between stable rotamer states (for example in eclipsed conformations), the final layer usually predicts a χ-angle closer to the canonical staggered rotamer states. As the χ values from the final layer are used to produce the AF2 structure, we hypothesised that the model may hold information about the distribution of possible χ_1_ states in the inner layers, followed by a switch to a single stable rotamer state for the final structure generation via the outer layer.

To test this hypothesis, we calculated χ_1_-angle averages from the UBQ DER ensemble and compared them with the χ_1_-angles from the AF2 structure module. We found that the χ_1_-angle averages from the inner layers correlated better (Pearson’s correlation coefficient *ρ*_*p*_=0.83) with the DER χ_1_ averages than χ_1_-angles from the last layer (*ρ*_*p*_=0.69) (Fig. 1g,h). A large fraction of χ_1_-angles have access to only one rotamer state, which may be confidently predicted by the final structure. We investigated whether the differences found between the inner and final layers were due to the subset of residues that populate multiple χ_1_ rotamer states based on the UBQ DER ensemble (Fig. 1i,j). For this subset of residues, we found an even greater difference in correlation between the inner (*ρ*_*p*_=0.80) and final (*ρ*_*p*_=0.62) layers. We made the same comparison for UBQ χ_2_-angles and found similar results (Fig. S2c,d), although with a smaller difference between the inner (*ρ*_*p*_=0.58) and final (*ρ*_*p*_=0.51) layers. Overall, these results suggest that the inner-layer χ-angle predictions may be sensitive to the underlying dihedral angle distribution; information that is discarded as the final layer is trained to produce a single side-chain configuration.

### Construction of AF2χ

Motivated by the correlation between χ-angle predictions from the inner layers of the AF2 structure module and DER ensemble averages, we developed a model that, given a protein sequence and backbone structure, uses these predictions to infer the underlying χ-angle distributions.

Bayesian/maximum entropy (BME) reweighting is a powerful approach to infer the distribution of structures underlying an ensemble-averaged observable (***Hummer and Köfinger, 2015***; ***Bottaro et al., 2020***). In our case, BME reweighting could be used to update a prior χ-angle distribution so the average is consistent with the χ-angle predicted by the inner layers of the AF2 structure module.

Here we introduce AF2χ, a model that uses BME reweighting to infer side-chain ensembles by integrating χ-angle predictions from the inner layers of the AF2 structure module with Bayesian priors based on the Top8000 rotamer library (***Hintze et al., 2016b***) and the AF2-predicted structure. AF2χ takes an input sequence and backbone structure, uses AF2 to extract information about side-chain χ_1_ and χ_2_ distributions, and generates an ensemble that predicts the structural heterogeneity of the side chains (Fig. 2).

**Figure 2.**
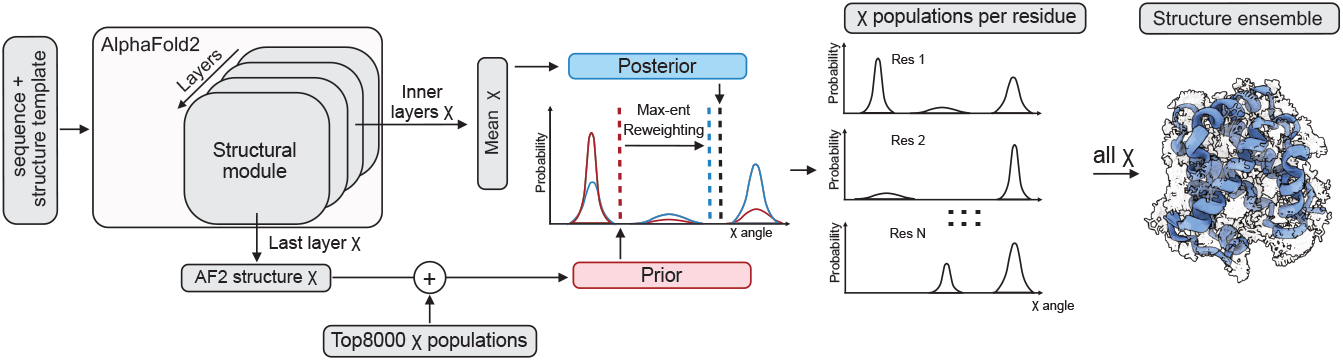
Schematic of AF2χ. Query sequence and structure template are used as input to AF2 from which the χ-angle values are extracted from the structure module. The prediction of the χ-angles from the last layer is combined with the Top8000 rotamer library to generate a prior distribution to be used during BME reweighting. The χ-angles from the inner layers are used as the posterior target ensemble average for BME reweighting. The output of BME is a collection of χ-angle distributions tailored to the query sequence and structure. Using AF2 and relaxation in a protein force field, the distributions are used to generate a conformational ensemble that reproduces side-chain structural heterogeneity.

#### Model parameterisation

Although AF2χ consists of several steps, the number of free parameters to set is relatively small; these are related to (i) the choice of parameters for the AF2 run, (ii) the creation of the prior distributions of χ-angles for input to BME, and (iii) the minimum number of structures to include in a generated structural ensemble. We again used UBQ as a pilot system to tune these free parameters, comparing model performance to HSP ensemble χ_1_ and χ_2_ distributions and experimental ^3^*J*-couplings from NMR reporting on χ_1_-angles.

We first tested different AF2 input parameters by generating a set of inner-layer χ-angles and calculating the correlation with the mean χ-angles from the UBQ DER ensemble distributions. Of the different setups tested (Fig. S3), we found two that had the best agreement with the DER ensemble averages: (i) AF2 with default parameters, namely full multiple sequence alignment, no template, and using the best ranked model, or (ii) AF2 in single-sequence mode, supported with a structure template, and using the best ranked model trained using templates. While we found that these setups gave similar accuracy, we decided to proceed with setup (ii) as the standard for AF2χ, as it allows for predictions based on any input structure template, making it possible to explore side-chain rotamer predictions for different backbone conformations of the query protein.

Next, we optimised how we generated the prior for BME reweighting as a weighted average of a Gaussian distribution around the AF2 structure χ-angle and the Top8000 distribution. By comparing the final BME distribution with ^3^*J*-couplings and the HSP ensemble χ_1_ and χ_2_ distributions (Fig. S4), given a mixing ratio, we optimised and selected the best mixing ratio between the two input distributions. We used the Jensen-Shannon (JS) divergence as the evaluation metric for the similarity to the HSP ensemble χ-distributions and the root mean square error (RMSE) for the similarity to ^3^*J*-couplings. We found that a mixture of the AF2-centered Gaussian and the Top8000 distribution gave results that were substantially better than using either individually. We set the mixing ratio to 15% Top8000 to 85% AF2 structure to balance the agreement with the HSP ensemble χ-distributions and ^3^*J*-couplings.

Next, we explored how the number of structures in the AF2χ structural ensemble affects the ability to accurately represent the dihedral distributions. As for the HSP ensembles, we used a bootstrap sampling procedure to generate AF2χ ensembles of different sizes and compared with ^3^*J*-couplings (Fig. S10). Consistent with our findings for HSP ensembles, we found that at least 20 structures were necessary to reproduce ^3^*J*-couplings. Based on these results, we generate AF2χ ensembles consisting of 100 structures.

The resulting AF2χ structural ensemble of UBQ has good agreement with the HSP ensemble (Fig. 3a,b; Tables S1, S2). We compared the results with baseline models such as the AF2 output structure, the Top8000 rotamer library, the DER ensemble, and MD simulations. We found that, for χ_1_ and χ_2_, AF2χ dihedral distributions have lower JS-divergence to HSP ensemble χ-angle distributions than the AF2 structure and Top8000. We also found that AF2χ dihedral distributions provide similar agreement with the HSP ensemble χ-angle distributions as MD simulations with CHARMM36m (***Huang et al., 2017***) and the UBQ DER ensemble. For residues that sample multiple χ_1_-angle free-energy wells in the HSP ensemble, we found that AF2χ provides better agreement with the HSP ensemble than CHARMM36m MD simulations, suggesting that AF2χ can accurately capture the structural heterogeneity of dynamic side chains.

**Figure 3.**
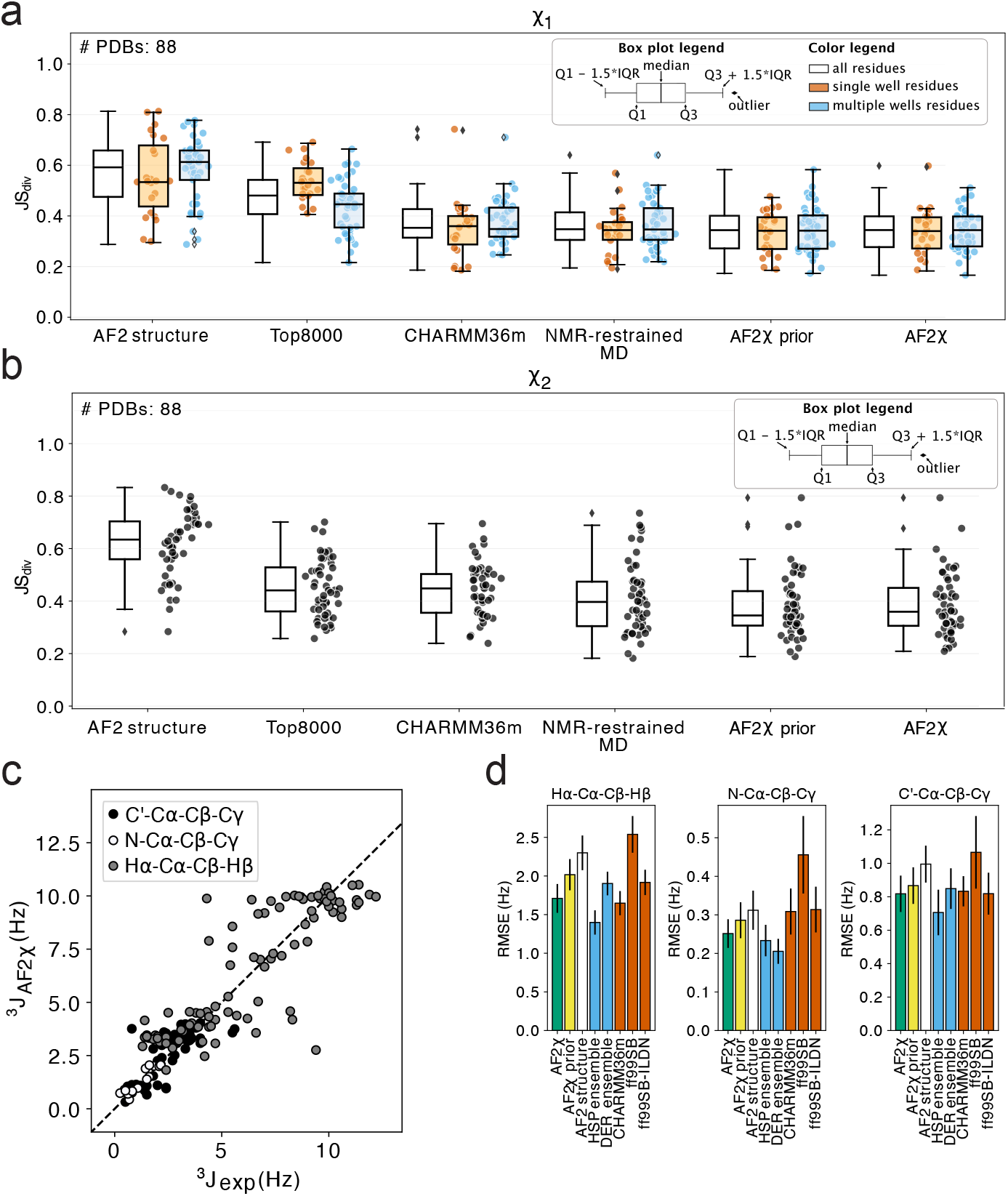
Benchmark of AF2χ predictions using HSP ensemble distributions and NMR ^3^*J*-couplings for UBQ. **(a**,**b)** Comparison of χ_1_ and χ_2_ distributions obtained with AF2χ to the UBQ HSP ensemble distributions using JS-divergence (JS_div_) as metric. χ distributions from other tested methods are reported for comparison. The summary statistics are reported with a box plot and the individual JS-divergence values are shown as dots. **(a**) χ_1_ JS-divergences to the HSP ensemble distributions for all UBQ residues (black), for single-well HSP residues (orange), and for multi-well HSP residues (light blue). **(b)** JS-divergences for χ_2_-angles to the HSP ensemble distributions for all residues. **(c)** Scatter plot with the ^3^*J*-couplings calculated from AF2χ structural ensembles (y-axis) and experimental values (x-axis). The different sets of UBQ ^3^*J*-couplings used are shown with different grey colours. **(d)** Bar plots reporting the root mean square error (RMSE) of ^3^*J*-couplings calculated from the AF2χ structural ensemble and other methods to experimental measurements. The RMSE values to experimental ^3^*J*-couplings reported are for: AF2χ (green), AF2χ prior distributions (yellow), AF2 structure (white), HSP and DER ensemble (blue), and MD simulations with different force fields (red). The black vertical line at the top of each bar shows the sample standard deviation of bootstrapped distributions.

The AF2χ ensemble of UBQ also has good agreement with NMR ^3^*J*-couplings (Fig. 3c,d; Table S3). Again, we compared AF2χ with several baseline models, including previously published ^3^*J*-couplings calculated from MD simulations with the Amber ff99SB (***Hornak et al., 2006***) and ff99SB-ILDN force fields (***Lindorff-Larsen et al., 2010***). We found that the AF2χ ensemble provides similar agreement with ^3^*J*-couplings as MD simulations, often reaching the accuracy of the ^3^*J*-couplings computed from the DER ensemble. We also found that the overall agreement improves as a result of the BME reweighting step for all three types of ^3^*J*-couplings, suggesting that the information from the inner layers of the AF2 structure module improves the accuracy of the predicted rotamer distributions (Fig. S5).

### Validation of AF2χ

We next validated AF2χ based on experimental data and MD simulations for independent protein systems.

#### Comparison with NMR ^3^*J*-couplings

To extend our validation of AF2χ, we calculated AF2χ dihedral angle distributions for three proteins with ^3^*J*-couplings from NMR reporting on χ_1_ dihedral angles: the B3 domain of Protein G (GB3), HEWL, and BPTI. As we had already constructed the HSP ensembles for HEWL and BPTI, we first compared the AF2χ dihedral distributions with the HSP ensemble distributions (Figs. S8 and S9) and found accuracy consistent with what we observed for UBQ; AF2χ matches the accuracy of CHARMM36m MD simulations for both χ_1_ and χ_2_ (Fig. S7,Tables S1, S2).

We then calculated ^3^*J*-couplings from the AF2χ ensembles of the three proteins, again comparing to predictions from AF2 structures, HSP ensembles, MD simulations with CHARMM36m, and previously published ^3^*J*-couplings from MD simulations with Amber ff99SB and ff99SB-ILDN (***Lindorff-Larsen et al., 2010***). We found that, overall, the AF2χ ensembles provide agreement with ^3^*J*-couplings comparable to the AF2 structure and atomistic MD simulations (Figs. 4 and S14; Table S3). Comparing the AF2χ priors with the final distributions, we found that the reweighting step of AF2χ slightly improves the agreement with experimental ^3^*J*-couplings for GB3, while it has no effect for HEWL and BPTI, suggesting that the information from the inner layers of the AF2 structure module in some cases can improve the accuracy of the χ_1_ distribution, as also observed for UBQ. Importantly, the BME reweighting step did not worsen the agreement for any of the proteins. The AF2χ prior provides an excellent description of the ^3^*J*-coupling data, suggesting that this very simple model produces reasonably accurate χ_1_ rotamer distributions. This is perhaps not surprising, as the AF2 structure, which the prior is largely based on, also provides reasonable agreement with the data.

**Figure 4.**
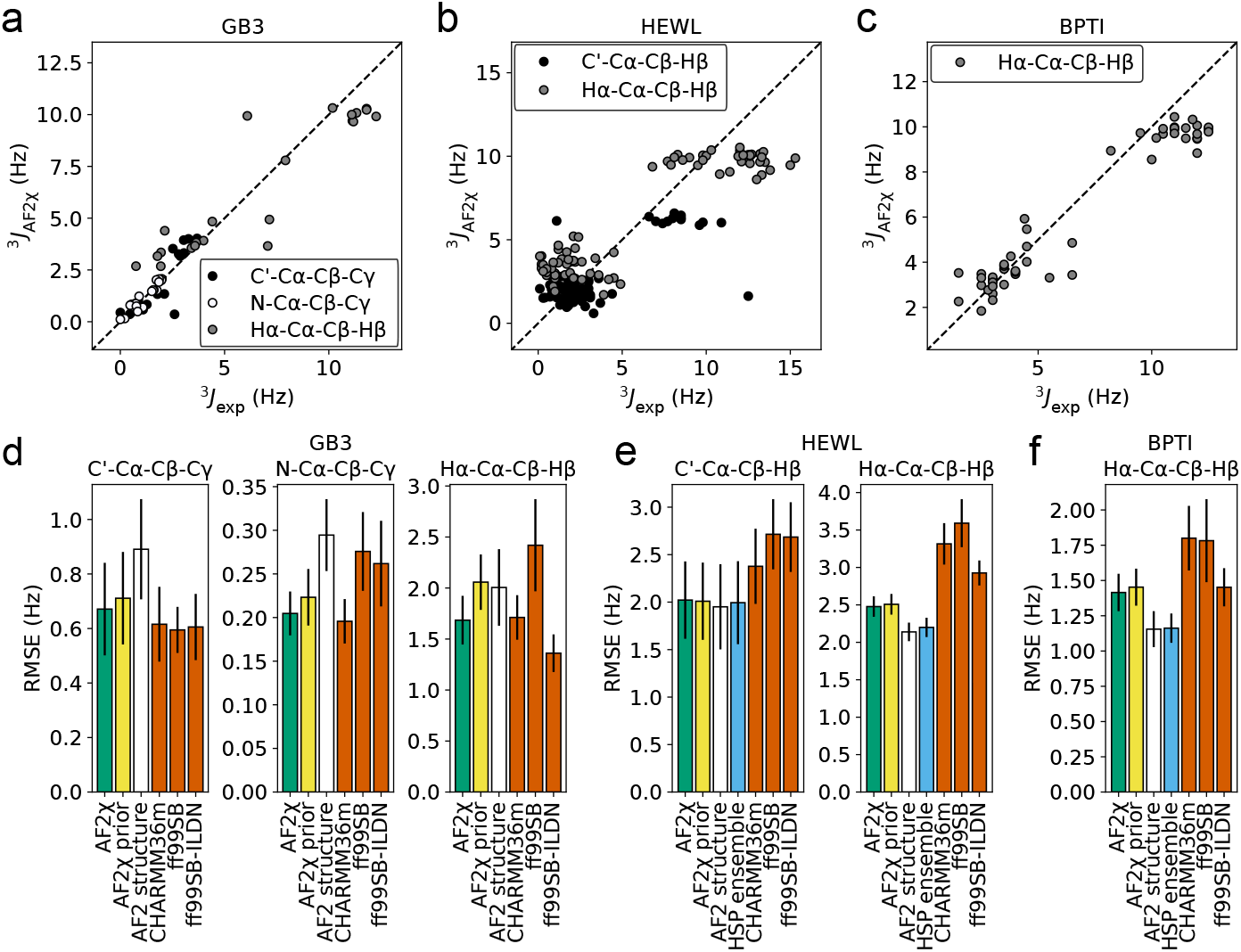
Validation of AF2χ structural ensembles using NMR ^3^*J*-couplings. **(a**,**b**,**c)** Scatter plots with ^3^*J*-couplings calculated from AF2χ structural ensembles (y-axis) and experimental values (x-axis) for the three validation proteins used: **(a)** GB3, **(b)** HEWL and **(c)** BPTI. For each protein, we show the different sets of ^3^*J*-couplings with a different grey colour. **(d**,**e**,**f)** Root mean square error (RMSE) of ^3^*J*-couplings calculated from AF2χ structural ensembles and other methods to experimental measurements for **(d)** GB3, **(e)** HEWL, and **(f)** BPTI. The RMSE values to the experimental ^3^*J*-couplings shown are for: AF2χ (green); AF2χ prior distributions (yellow), AF2 structure (white), HSP ensemble (where available) (light blue), and MD simulations with different force fields (red). The black vertical line at the top of each bar shows the sample standard deviation of bootstrapped distributions.

#### Comparison with NMR side-chain methyl order parameters

To further validate the prediction of side-chain dynamics by the AF2χ ensembles, we compared with experimental side-chain methyl 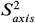 order parameters from NMR for the third fnIII domain from tenascin (TNfn3), the tenth fnIII domain from fibronectin (FNfn10), adipocyte lipid binding protein (A-LBP), muscle fatty acid binding protein (M-FABP), A3D, and Fyn SH3. Methyl 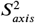 order parameters depend on the motions of the side-chain methyl C-C bond vector, thereby probing the conformational heterogeneity of side chains, and can be estimated from ensembles or simulations using a model that depends on ensemble-averaging of bond vector components.

We generated AF2χ structural ensembles for the six proteins and calculated 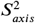 order parameters for the methyl groups corresponding to the experimental data. For comparison, we also performed MD simulations of the six proteins using the CHARMM36m force field and calculated 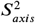 order parameters from our simulations (Figs. 5 and S15; Table S4). Overall, we found that the AF2χ ensembles and MD simulations provide comparable correlations with the experimental 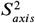 data. For A3D, AF2χ gave a higher correlation with the experimental data than MD, and vice versa for Fyn SH3. We also calculated 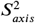 order parameters from structural ensembles produced with the AF2χ prior. We found that the BME reweighting step of AF2χ improves the correlation with the experimental 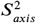 order parameters for four out of six proteins, and does not worsen the agreement for any protein (Fig. 5b), again suggesting that the information from the inner layers of the AF2 structure module improves the accuracy of the predicted side-chain ensemble.

**Figure 5.**
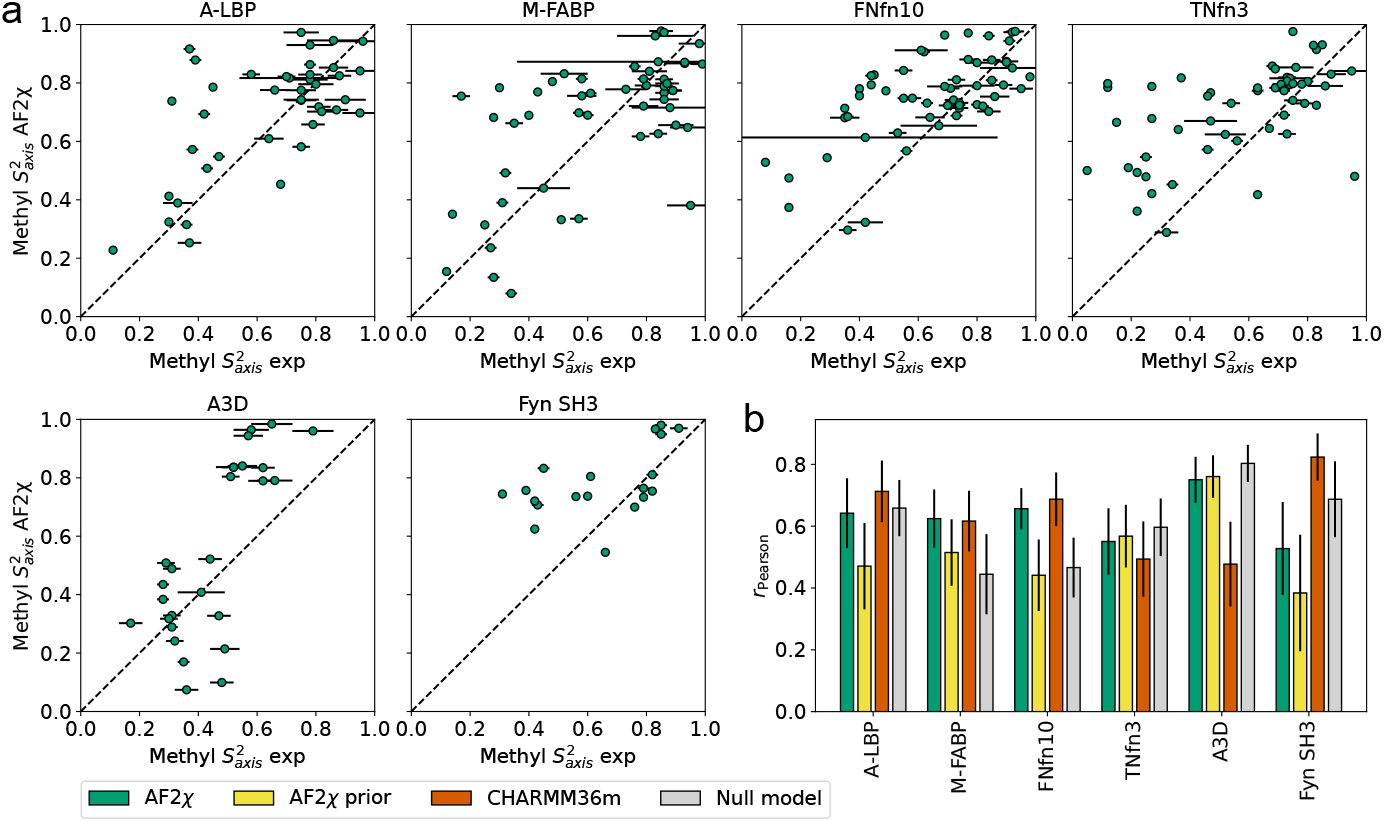
Validation of AF2χ structural ensembles using NMR methyl 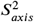 order parameters. (a) Comparison of experimental methyl 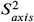 (x-axis) and those calculated from AF2χ structural ensembles (y-axis) for six benchmark proteins (A-LBP, M-FABP, FNfn10, TNfn3, A3D, and Fyn SH3). Experimental errors are shown as black lines for each point. **(b)** Pearson correlation coefficient (*r*_*pearson*_) between the methyl 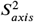 order parameters calculated from the AF2χ structural ensemble (green) and the other methods used as benchmarks: AF2χ prior (yellow), null model (grey), and MD simulations with CHARMM36m (red). The black vertical line at the top of each bar shows the sample standard deviation of bootstrapped distributions.

In general, AF2χ slightly overestimates 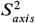 values, suggesting that side-chain mobility may be slightly underestimated. As NMR 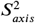 order parameters capture motions on time scales shorter than the global tumbling of the protein, this suggests that the mobility of side-chains in AF2χ corresponds to motions on the ps-to-ns time scale. This is consistent with what has been observed for HSP ensembles (***Best et al., 2006***), and other studies that suggest that much (***Chou et al., 2003***; ***Lindorff-Larsen et al., 2005b***; ***Xiang et al., 2021***; ***Hoffmann et al., 2021***), but not all (***Fares et al., 2009***; ***Shaw et al., 2010***; ***Long et al., 2011***; ***Genheden et al., 2014***; ***Smith et al., 2015***), side-chain mobility occurs within this time scale.

Side-chain methyl 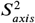-values are correlated (***Mittermaier et al., 1999***; ***Best et al., 2004***) with residue and methyl type (e.g. Ile C*γ*), so model predictions should be compared with a null model that predicts the 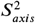 value as a function of methyl type without taking into account the structural context. We calculated the average experimental 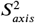 across our set of proteins for each methyl type and used this as a null model to predict 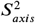. For two proteins, M-FABP and A-LBP, we found that the AF2χ ensembles and MD simulations gave a higher correlation with experimental 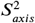 values than the null model, while the MD simulations also gave a higher correlation than the null model for Fyn SH3. These results suggest that, although the AF2χ ensembles provide agreement with experimental 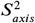 values comparable to MD simulations, neither of the models accurately capture site-specific relative differences in 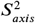 for specific methyl types.

#### Comparison with ATLAS MD simulation database

To investigate how well AF2χ agrees with side-chain rotamer distributions predicted by atomistic MD simulations, we generated AF2χ dihedral distributions for the 1390 proteins in the ATLAS MD database (***Meersche et al., 2024***). ATLAS contains 300 ns (three simulations each 100 ns long) MD simulations with the CHARMM36m force field for a diverse set of proteins. We calculated χ_1_ and χ_2_ dihedral angle distributions from the ATLAS simulations and evaluated the agreement with AF2χ based on the JS-divergence (Fig. 6a). As baseline models, we used the Top8000 rotamer library distributions and AF2 structures of the 1390 proteins. For both χ_1_ and χ_2_, we found that AF2χ is in reasonable agreement with MD simulations, with the median JS-divergence across all proteins under 0.4. For χ_1_, AF2χ provides better agreement with the simulations than both the AF2 structure and Top8000. For χ_2_, AF2χ also provides better agreement than the AF2 structure, but comparable agreement with Top8000. These results suggest that AF2χ contains structure-based information about the side-chain rotamer distributions that better reflects the solution ensemble than the static AF2 structure.

**Figure 6.**
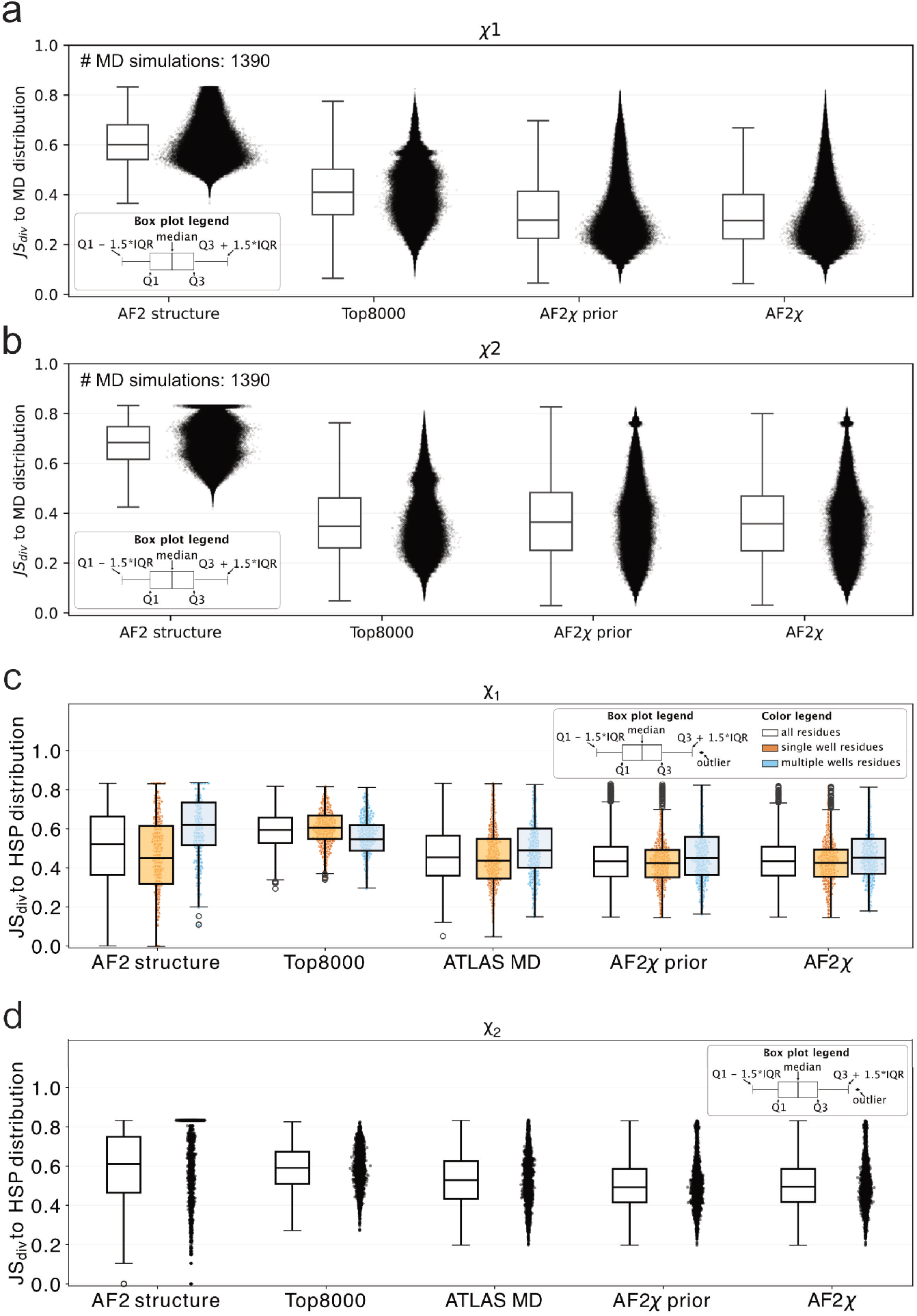
Large-scale benchmark of the AF2χ structural ensemble against the ATLAS MD database. **(a**,**b)** JS-divergence values (y-axis) for **(a)** χ_1_ and **(b)** χ_2_ distributions from AF2χ to the distributions for 1390 proteins obtained from ATLAS MD simulations. We report a box plot summarising the JS divergence statistics (left) and all the JS data points (right) for each method: AF2 structure (left), Top8000 (middle-left), AF2χ prior (middle-right), and AF2χ (right). **(c)** JS-divergence values for χ_1_-angle distributions to HSP ensembles of 15 proteins. The left box plot (white) shows the JS-divergence statistics for all residues in the 15 proteins, the middle box plot (orange) for the HSP single-well residues, and the right box plot (light blue) for the HSP multiple-well residues. The individual JS divergences are shown as dots below the single- and multiple-well box plots. **(d)** JS-divergence values for χ_2_-angle distributions to HSP ensembles of 15 proteins. The box- and scatter-plots show the JS-divergence values for all residues of the 15 proteins for the different methods tested.

#### Comparison with HSP ensembles for ATLAS proteins

For 15 monomeric proteins in the ATLAS database, we found 10 or more high sequence-similarity structures in the PDB and used these to generate HSP ensembles. Given the accuracy of HSP in mimicking solution state dynamics, this enabled us to compare the accuracy of the ATLAS MD simulations with ensembles generated by AF2χ. We compared the AF2χ (χ_1_ and χ_2_) distributions to the HSP ensemble distributions, using as baseline models the AF2 structure, the Top8000 rotamer library, and the ATLAS MD simulations (Fig. 6b). Consistent with our other results, AF2χ, MD simulations, and the AF2 structure gave similar JS-divergences to the HSP ensemble distributions, and better agreement than Top8000. Again, we classified residues into those that populate a single χ_1_-angle free energy well or multiple free-energy wells based on the HSP ensemble. We found that, for residues that populate multiple χ_1_ free-energy wells, AF2χ provides substantially better agreement with the HSP ensemble than the AF2 structure and slightly better agreement than the ATLAS MD simulations. For residues with a single χ_1_ free energy well, the AF2 structure, ATLAS MD simulations, and AF2χ provide a similar median agreement with the HSP ensemble, while the distribution of JS divergences is broader for the AF2 structure because it only populates a single angle. Finally, we observed (Fig. S12) that AF2χaccurately predicts HSP ensemble distributions for proteins up to 1000 residues, and that its accuracy does not decrease with protein size. Overall, these results show that AF2χ dihedral distributions are as accurate as those extracted from MD trajectories in recovering HSP ensemble distributions.

#### Computational cost of AF2χ

While providing similar accuracy as MD simulations in our tests, AF2χ represents a computational speedup in the generation of side-chain dihedral distributions and structural ensembles. We compared the computational cost of AF2χ runs with MD simulations for all the proteins in the ATLAS MD database (Fig. S13). We found that AF2χ predicts rotamer distributions at a speed of 0.25 seconds per residue, even for large protein systems, whereas the ATLAS MD simulations scale roughly linearly with protein size to about 1 day per 100 residues simulated. When we also include the generation of the structural ensembles, we found that AF2χ remains up to three orders of magnitude faster than MD simulations, with computational times scaling linearly with protein size to about 10 minutes per 100 residues for a 100-structure ensemble.

### Comparison with generative models for protein ensembles

A number of methods have been developed to predict side-chain conformations (***Misiura et al., 2022***; ***Visani et al., 2023***; ***Mukhopadhyay et al., 2023***; ***Guerois et al., 2002***; ***Eyal et al., 2004***; ***Liang et al., 2011a***; ***Hartmann et al., 2007***; ***Lu et al., 2008***; ***Liang et al., 2011b***; ***Miao et al., 2011***; ***Xiang et al., 2007***; ***Peterson et al., 2014***; ***Krivov et al., 2009***; ***Yang et al., 2002***; ***Hwang and Liao, 1995***; ***Peterson et al., 2004***); however, with notable exceptions (***Kussell et al., 2001***; ***Gautier and Tufféry, 2003***; ***Hu and Kuhlman, 2006***; ***DuBay et al., 2011***; ***Bhowmick and Head-Gordon, 2015***), most of these methods aim to recover the dominant side-chain conformation, rather than the conformational heterogeneity of the side chain. As AF2χ aims to generate distributions of side-chain structures, we decided to compare to recently developed generative models that focus on recovering the structural dynamics of the backbone and side chains. We thus benchmarked AF2χ against side-chain rotamer distributions generated from: (i) structural ensembles created using AF2 with systematic residue dropout, a technique that has been shown to facilitate the exploration of alternative protein conformations (***Kalakoti et al., 2025***), (ii) MDgen (***Jing et al., 2024b***), (iii) SeqDance (***Hou and Shen, 2024***), (iv) BioEmu + Hpacker (***Visani et al., 2023***; ***Lewis et al., 2024***) and (v) aSAMt (***Janson et al., 2025***). We note here that these models were developed and benchmarked to predict conformational dynamics mostly at the backbone level, though to some extent also for side chains.

We tested the performance for UBQ, HEWL, and BPTI, as these proteins had both HSP ensembles and ^3^*J*-coupling data available. We found that for all three proteins, AF2χ generates distributions in better agreement with HSP ensemble distributions than the other models (Fig. S11a,b,c; Table S5). Of the other tested generative models, MDgen, BioEmu + Hpacker, and aSAMt have the best agreement with HSP ensemble distributions. AF2 dropout has improved agreement with HSP ensembles compared with the AF2 structure; however, it does not achieve comparable accuracy to the other models on residues with multiple χ_1_ free-energy wells. SeqDance has the worst agreement of the models tested, with distributions very different from the HSP ensemble distributions. The lower accuracy of SeqDance might be related to the use of sequence as the only input for prediction of χ-angle distributions and a sampling binning step of 30 degrees.

Given the overall good performance of MDgen, BioEmu + Hpacker, and aSAMt when comparing with HSP ensembles, we also tested these models for agreement with ^3^*J*-couplings (Fig. S11d,e,f; Table S6). We found that, of the four models, AF2χ ensembles have the best overall agreement with ^3^*J*-couplings.

## Conclusions

Here we have introduced AF2χ, a model to predict side-chain rotamer distributions and generate side-chain structural ensembles for an input sequence and structure. The foundation of AF2χ is the ability of AF2 to predict χ-angles. Recent results (***Maisuradze et al., 2025***) have shown that AF2 is accurate at predicting the main state of rotamers, with greater accuracy for χ-angles closer to the backbone. Our investigation extends these results and shows that AF2 has also learned information about side-chain structural heterogeneity in the form of averaged χ-angle predictions in the inner layers of the structure module. This information has likely been learnt by AF2 because torsional losses are evaluated only on the last-layer values, and because of the extensive training on a large number of amino acids in different physico-chemical environments, which—similar to HSP ensembles—may represent protein dynamics and enable AlphaFold in some sense to learn a physical potential (***Roney and Ovchinnikov, 2022***). These observations also suggest that it may be possible to train a model more directly to predict side-chain dynamics, for example by targetting experimental HSP ensembles, or by targetting conformational heterogeneity across predictions of related proteins (***Lewis et al., 2024***).

HSP ensembles are an approach based on overlaying experimental protein structures with high sequence-similarity to predict structural dynamics (***Best et al., 2006***). In the original work, HSP ensembles were validated using mainly NMR order parameters, while a direct comparison with ^3^*J*-couplings was only done for HIV-1 protease. Here, we show that HSP ensembles of UBQ, HEWL, and BPTI also reproduce ^3^*J*-couplings reporting on χ_1_ with high accuracy.

We observed that the AF2χ prior provides reasonable agreement with NMR ^3^*J*-couplings, HSP ensembles, and MD simulations, suggesting that the structural heterogeneity of protein side chains in some cases can be reasonably captured by predicting dihedral angle distributions with a simple combination of the AF2 structure and Top8000 rotamer library. This simple model performs relatively well and is therefore a useful null model to benchmark methods for predicting side-chain ensembles or dihedral angle distributions.

Generative ML models models for predicting structural ensembles of proteins (***Stein and Mchaourab, 2022***; ***del Alamo et al., 2022***; ***Jing et al., 2023, 2024a***,b; ***da Silva et al., 2024***; ***Lewis et al., 2024***; ***Wayment-Steele et al., 2024***; ***Qiao et al., 2024***; ***Zheng et al., 2024***; ***Lu et al., 2024***) are often benchmarked by comparing with MD simulations, which are limited in accuracy by the quality of the force fields and sampling, or by comparing with experimentally determined structures of different conformations, which lack quantitative information about the associated free energies. Therefore, comparison of predicted protein ensembles with solution NMR data and HSP ensembles is a useful additional benchmark, as these more directly report on equilibrium structural heterogeneity. We believe that the dataset collected from the literature in this work will be useful for future model development, in particular for models that predict protein side-chain dynamics.

As AF2χ initially models the dihedral angle distributions of each residue separately, correlations between side-chain conformations are not captured in this step of the model. In principle, correlations between side-chain conformations could be captured in the structure generation step of AF2χ, as structures with clashes are rejected; in practice we find that the acceptance rate of structures is close to 100%. This suggests that any remaining steric clashes may be resolved when the backbone structure is relaxed, in line with previous observations that small backbone movements help decouple side-chain motions (***Davis et al., 2006***; ***DuBay et al., 2011***). These observations are also consistent with previous observations that dynamics of neighbouring side chains are not strongly correlated (***Best et al., 2005***; ***Hoffmann et al., 2021***).

The ability to predict the flexibility of side chains in proteins is important for many tasks, from structure prediction to computational design of proteins and ligands. Our method generates backbone-specific dihedral distributions and provides an efficient and relatively accurate description of the side-chain free energy landscapes. In our study we focused particularly on an extensive validation of the predictions of dihedral distributions and the quality of the generated structural ensemble.

There are many uses for the dihedral distributions that AF2χ outputs, for example as a starting point to study differences in side chain conformations for different conformational states (***Tuttle et al., 2013***) or to help interpret experimental data probing how amino acid substitutions may propagate effects to neighbouring side-chains (***Fraser et al., 2009***). Furthermore, structural ensembles can be used as input to design tools to improve the quality of protein representation (***Beglov et al., 2012***). Additionally, AF2χ ensembles could have applications in ensemble docking, conformational entropy calculations, and experimental modelling, such as fitting cryo-EM and X-ray density maps or modelling fluorescent dyes or spin-labels.

In summary, AF2χ makes it possible to extract information about protein side-chain rotamer distributions from the AF2 structure module and generate structural ensembles of side chains. We show that AF2χ ensembles reproduce NMR ^3^*J*-couplings, NMR methyl 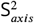 order parameters, and HSP ensembles with accuracy similar to atomistic MD simulations, while AF2χ generates side-chain rotamer distributions three orders of magnitude faster than MD simulations. Given the speed and accuracy of AF2χ side-chain ensemble predictions, we envisage that it will be a powerful tool for protein design, ligand docking, and interpretation of biophysical experiments.

## Methods

### AF2χ model

#### AF2 run setup

AF2χ runs AF2 with LocalColabFold v.1.5.5 (***Mirdita et al., 2022***) and can be run on both monomeric and oligomeric structures, using either the AF2 `alphafold2` or `alphafold2_multimer_v3` models; no flag is required in the input to switch between the two models. We have found two optimal AF2 setups to use with AF2χ:

- AF2χ with standard AF2 setup, where we use the full MSA and no structural templates as input to the model. In this setup we used predictions only from three of the five models (model 1.2.1,1.2.2,1.2.3) as they were trained in the same regime. This setup is preferred when the target structure is the native state of the protein but the structure is unknown. The AF2χ standard run is carried out in localcolabfold using the flags: `-msa-mode mmseqs2_uniref_env` to use the full MSA processing and `–model-order 3,4,5` to restrict use to the three selected models.
- AF2χ with decoy strategy (***Roney and Ovchinnikov, 2022***). The decoy strategy uses the query sequence as input to AF2 supported by a custom structure template and the complete removal of the MSA. We implemented the decoy strategy as the main AF2 setup for AF2χ to enable sampling of dihedral distributions and generation of structural ensemble around any input template structure. The AF2χ decoy strategy is run in LocalColabFold using the flags: `–msa-mode single_sequence` to remove the MSA processing, `–template` and `–custom-template` to enable the custom template structure, and `–model-order 1,2` to use only the two models (1.1.1 and 1.1.2) that were trained to use information from template structures.

#### Extraction of AF2 χ-angles

AF2χ uses the structure module of AF2 to extract χ angle information. In particular, the multi-rigid side-chain unit (Algorithm 20, lines 11-14, AF2 Supplementary Information) generates torsion angles for the backbone and side-chain. For each iteration, the structure module is run eight times, each time generating a complete prediction for all side-chain torsion angles. In AF2χ we use the χ-angle predictions from the last iteration. We use the average of the predictions from the second to the seventh structure module run (here inner layers) to evaluate the target ensemble average for our BME reweighting. We also use the inner layers to estimate the error (standard deviation) of the target ensemble mean. The predictions from the last structure module run (last layer) are used to generate the AF2 output structure without any further correction, thus we use these values instead of the χ-angles extracted from the final structure to generate our Bayesian priors.

#### Priors

As Bayesian priors for AF2χ, we constructed dihedral angle distributions for each χ-angle by calculating a weighted sum of a Gaussian distribution centered on the χ-angle of the AF2 structure (extracted from the last layer of the AF2 structural module) and the Top8000 rotamer library distribution. The AF2χ prior is given by:

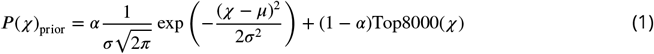

where *µ* is set to the χ-angle calculated from the AF2 structure, and we determined amino acid- and χ-angle-specific values for *σ* by fitting Gaussians to the main populations of the Top8000 distributions and averaging the resulting *σ*-values (e.g. averaging the fit of three Gaussians for Phe χ_1_ and two for Phe χ_2_). To determine the value of *α*, we scanned values from 0 to 1 in steps of 0.05 and calculated the agreement with NMR ^3^*J*-couplings and HSP ensemble distributions for UBQ. For each value of *α*, we also ran the full AF2χ calculation and evaluated the agreement with the data. Based on this analysis, we select *α*=0.85.

The Top8000 rotamer library consists of, for each amino acid type, a multidimensional dihedral angle distribution that is a function of all χ-angles. To obtain independent distributions for each χ-angle, we integrated the probabilities over the remaining χ-angles. Finally, we rebinned the Top8000 dihedral angle distributions to 36 bins.

#### BME reweighting

AF2χ uses BME reweighting to update the prior dihedral angle distribution to fit the inner-layer angle prediction as a target ensemble average. We fit a set of *n* weights (*w*_1_, …, *w*_*n*_), in this case the probabilities of the 36 dihedral angle bins, by minimizing the following loss function:

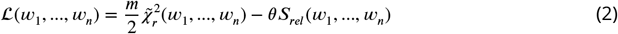

where *m* is the number of target data points (in this case *m* = 1) and *θ* is a global scaling parameter that balances the confidence in the prior versus the target data. For each χ-angle, we determine *θ* by scanning 20 values from 1 to 10.000 evenly spaced on a log scale and selecting the highest value that results in 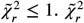 is the reduced chi-square, which quantifies the agreement between the dihedral angle distribution and the inner-layer prediction and is given by:

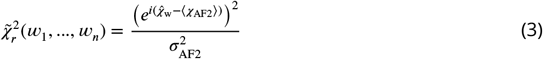

where ⟨*x*_AF2_⟩ and *σ*_AF2_ are the average and associated standard deviation of dihedral angles from the inner layers of the AF2 structure module. 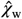 is the circularly averaged dihedral angle calculated from the dihedral distributions using:

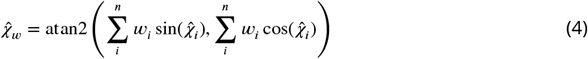

where 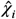 are the discrete *χ*-angle values from the distribution and *w*_*i*_ the corresponding weights. *s*_*rel*_ is the relative Shannon entropy between the prior (denoted by *w*^0^) and posterior weights (*w*):

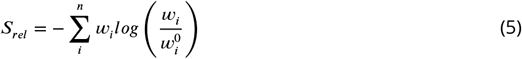

We minimised equation 2 using SLSQP minimization in SciPy (***Virtanen et al., 2020***) with a constraint on the sum of the weights:

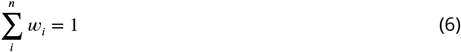

and bounds on the weights: 0 ≤ *w* ≤ 1.

#### Generating Structural ensembles

We used the AF2 internal function and classes to generate our structural ensemble. The protocol is divided into: (1) structure generation, (2) structure relaxation, (3) checking for clashes, and (4) checking for agreement with the AF2 structure.

1. To generate a structure after generating dihedral distributions, we create spatial quaternions from the backbone torsion angles of the last layer of the structural module (the code implementation here differs slightly between monomers and oligomers). We then use the AF2χ dihedral distributions to sample χ_1_ and χ_2_ values for each residue and add them to the quaternion representation. We convert the quaternions to atomic 3D coordinates and use the supporting utilities function of AF2 to generate a coordinate file.
2. The structure is then relaxed using the standard AF2 protocol, which consists of energy minimization with the Amber ff99SB force field (***Hornak et al., 2006***) using OpenMM (7.7.0) (***Eastman et al., 2017***). To ensure that the angles sampled from the AF2χ distributions are retained in the final structure, we add periodic torsion restraints for all χ_1_ and χ_2_ angles with a force constant of 10000 kJ mol^−1^. As a standard in AF2, harmonic position restraints with a force constant of 10.0 kcal mol^−1^ Å^−2^ are added for all heavy atoms; we lowered this force constant to 0.5 kcal mol^−1^ Å^−2^ to allow for more movement of the backbone to accommodate clashing side chains.
3. We then check the relaxed structure for atomic clashes by calculating the atomic distance for all atoms pairs in different residues and compare to a distance threshold:

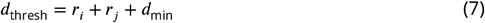

where 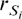 and 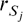 are the atomic radii for the two atoms involved and *d*_tmin_ is a user-defined minimum distance between the two atoms. We used *d*_thresh_ = 0.65 Å for all our runs. We discard any structure that still has atomic clashes after relaxation.
4. Finally, we calculate the root mean square deviation (RMSD) from the final AF2 structure to verify that the relaxed structure remained close to the input. We use a relative RMSD threshold (RMSD_thresh_) that scales with the size of the query protein (***Carugo and Pongor, 2001***), defined as:

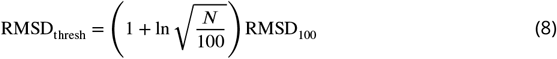

where *N* is the size of the protein and RMSD_100_ is a selected reference threshold for a protein of length 100 residues. If the RMSD of the relaxed structure is greater than the threshold, the structure is discarded. In our work we chose RMSD_100_ to be 1.0Å, so that the RMSD threshold for example is 0.65Å for a protein of 50 residues and 2.15Å for a protein of 1000 residues.

## NMR ^3^*J*-couplings calculations

We compared with the set of experimental NMR ^3^*J*-couplings for UBQ, HEWL, BPTI and GB3 previously collected in ***Lindorff-Larsen et al***. (***2010***). We generated AF2χ ensembles using the following structures as templates: UBQ: PDB 1UBI (***Ramage et al., 1994***), GB3: PDB 1P7E (***Ulmer et al., 2003***), HEWL: PDB 6LYT (***Young et al., 1993***), and BPTI: PDB 5PTI (***Wlodawer et al., 1984***).

We compared with H*α*-C*α*-C*β*–H*β*1,2 ^3^*J*-couplings for all four proteins (***Smith et al., 1991***; ***Berndt et al., 1992***; ***Schwalbe et al., 2001***; ***Miclet et al., 2005***), N-C*α*-C*β*-C*γ* ^3^*J*-couplings for GB3 and UBQ (***Chou et al., 2003***), C’-C*α*-C*β*-C*γ* ^3^*J*-couplings for GB3 (***Chou et al., 2003***) and UBQ (***Hu and Bax, 1997***), and C’-C*α*-C*β*-H*β*1,2 ^3^*J*-couplings for HEWL (***Grimshaw, 1999***). ^3^*J*-couplings were calculated from χ_1_ dihedral angles of structures using the Karplus relationship:

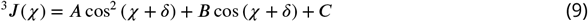

where *χ* is the dihedral angle and *A, B, C*, and *δ* are parameters specific to the residue and dihedral angle type (***Karplus, 1963***). We used Karplus parameters from ***Chou et al***. (***2003***) for Val, Ile, and Thr N/C’-C*α*-C*β*-C*γ* ^3^*J*-couplings and Karplus parameters from ***Pérez et al. (2001)*** for all other ^3^*J*-couplings. We also compared the experimental ^3^*J*-couplings with values reported in ***Lindorff-Larsen et al***. (***2010***) for MD simulations with Amber ff99SB and ff99SB-ILDN. The errors on the RMSE values between experimental and ensemble ^3^*J*-couplings were calculated by bootstrapping the set of ^3^*J*-couplings using SciPy (1.13.0).

## NMR methyl 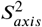 order parameters calculations

We compared with experimental NMR methyl 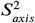 order parameters for Fyn SH3 (***Mittermaier et al., 2003***), TNfn3, FNfn10 (***Best et al., 2004***), A-LBP, M-FABP (***Constantine et al., 1998***), and A3D (***Walsh et al., 2001***). We generated AF2χ ensembles using the following structures as templates: A-LBP: PDB 2HNX (***Marr et al., 2006***), M-FABP: PDB 1HMT (***Young et al., 1994***), FNfn10: PDB 1FNF (***Leahy et al., 1996***), TNfn3: PDB 1TEN (***Leahy et al., 1992***), A3D: PDB 2A3D (***Walsh et al., 1999***), and Fyn SH3: PDB 1SHF (***Noble et al., 1993***).

We calculated 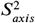 from structural ensembles as:

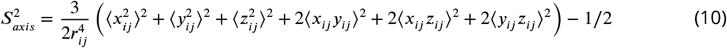

where *x*_*ij*_, *γ*_*ij*_, and *z*_*ij*_ are the components of the interatomic bond vector between carbon *i* (bonded to a methyl group) and carbon *J* (in a methyl group), *r*_*ij*_ is the length of the C-C bond, and the angle brackets denote averaging over the ensemble structures (***Henry and Szabo, 1985***; ***Best and Vendruscolo, 2004***). To exclude the contribution of motions on time scales longer than the rotational correlation time, we split our MD simulations into blocks of 10 ns (i.e. 100 blocks for our 1 *µ*s simulations), calculated 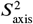 for each block, and averaged the values to obtain the final 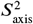. Structures were superposed to the ensemble-averaged structure prior to 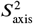 calculations. The errors on the Pearson correlation coefficients (*r*_Pearson_) between experimental and ensemble 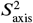 values were calculated by bootstrapping the set of 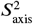 values using SciPy (1.13.0).

## Molecular dynamics simulations

We ran MD simulations of the following proteins using the corresponding PDB structure as the starting structure: UBQ (PDB 1UBI (***Ramage et al., 1994***)), GB3 (PDB 1P7E (***Ulmer et al., 2003***)), HEWL (PDB 6LYT (***Young et al., 1993***)), BPTI (PDB 5PTI (***Wlodawer et al., 1984***)), A-LBP (PDB 2HNX (***Marr et al., 2006***)), M-FABP (PDB 1HMT (***Young et al., 1994***)), FNfn10 (PDB 1FNF (***Leahy et al., 1996***)), TNfn3 (PDB 1TEN (***Leahy et al., 1992***)), A3D (PDB 2A3D (***Walsh et al., 1999***)), and Fyn SH3 (PDB 1SHF (***Noble et al., 1993***)). We ran simulations with the CHARMM36m protein force field and CHARMM-modified TIP3P water model (***Huang et al., 2017***) using Gromacs 2021.1 (***Abraham et al., 2015***). We placed each protein structure in a dodecahedral box, with the box size set to have a minimum distance of 1.1 nm between the protein and the box edge. We solvated the system and added 150 mM NaCl with additional Na^+^ or Cl^−^ ions to neutralise the system.

Simulations were performed using the leap-frog integrator with a 2 fs time step. We used a Verlet list cut-off scheme for non-bonded interactions. We used a 1.2 nm cut-off for van der Waals interactions with a switching function starting at 1.0 nm. We used Particle Mesh Ewald (PME) for electrostatic interactions with a real-space cutoff of 1.2 nm, a Fourier grid spacing of 0.12 nm, and fourth order cubic interpolation. H-bonds were constrained with LINCS (***Hess et al., 1997***). We used the velocity rescaling thermostat (V-rescale) at 300 K with a 0.1 ps time coupling constant (***Bussi et al., 2007***) and the Parrinello-Rahman barostat with a pressure of 1.0 bar, a 2 ps time coupling constant, and an isothermal compressibility of 4.5 × 10^−5^ bar^−1^ (***Parrinello and Rahman, 1981***). Production simulations were run for 1 *µ*s with protein coordinates saved every 100 ps.

Before production simulations, we performed steepest descent energy minimization with a step size of 0.01 nm until a force of <1000.0 kJ mol^−1^ nm^−1^ was reached. We then generated velocities sampled from the Maxwell-Boltzmann distribution at 300 K and performed a 3-step equilibration protocol: (1) 2 ns NVT simulation with a 1 fs time step with harmonic position restraints on all protein heavy atoms with a force constant of 1000 kJ mol^−1^ nm^−2^, (2) 2 ns NPT simulation with a 1 fs time step with the same position restraints, (3) 10 ns NPT simulation with a 2 fs time step. NPT equilibration simulations were performed with the stochastic cell rescaling (C-rescale) barostat with a pressure of 1.0 bar, a 5 ps time coupling constant, and and an isothermal compressibility of 4.5 × 10^−5^ (***Bernetti and Bussi, 2020***).

## ATLAS MD calculations

We downloaded MD simulation trajectories for the 1390 proteins in the ATLAS MD database (***Meersche et al., 2024***) and calculated χ_1_ and χ_2_ dihedral angle distributions from the trajectories using MDTraj (1.9.9) (***McGibbon et al., 2015***). For each protein, we also downloaded the starting structure for the MD simulation and used this as a template for AFχ to predict χ_1_ and χ_2_ dihedral angle distributions. We compared the χ_1_ and χ_2_ dihedral angle distributions from AFχ, Top8000, and the AF2 structure with the ATLAS MD distributions using the JS divergence over probability histograms with 36 bins.

## Generative models for protein ensembles

### AF2 dropout

We introduced dropout in AF2 by using the corresponding option available in localColabFold 1.5.5, implemented with the flag `-use-dropout`, which stochastically drops some of the weights in the model. We used the dropout option in combination with both AF2 inference setups (standard and decoy). We found no differences between the two. For each protein, we generated an ensemble of 100 structures.

### Seqdance

We used SeqDance_35M (***Hou and Shen, 2024***) built over ESM2 to infer the side chain torsion angles from the output features, obtained by running a single forward sampling. As SeqDance creates χ-angle distributions with a 30° binning interval, we evaluated the JS divergence on re-binned HSP distributions over twelve bins, following the SeqDance binning strategy.

### BioEMu and HPacker

We used BioEmu v.0.1.6 (***Lewis et al., 2024***) to generate a set of backbone conformations for each of the three benchmark proteins using `bioemu.sample` with the MSA generated internally with ColabFold. We then examined that the generated backbones were close to our reference structure and discarded them if their RMSD from the reference was greater than 2Å. We aimed to generate 100 structures for each protein. We then ran HPacker (***Visani et al., 2023***) with the `bioemu.relax` prompt using the backbone structures generated by Bioemu. We used the `–no-md-equil` flag to avoid performing additional MD relaxations that would change the produced χ angles.

### aSAMt

We used aSAMt (***Janson et al., 2025***) to generate structural ensembles for our benchmark proteins. We sampled at a temperature of 298K, with a batch size of 8. We examined whether the ensemble structures were close to our reference structure, and discarded them if their RMSD from the reference was greater than 2Å. We generated an ensemble of 100 structures for each protein.

## Data and code availability

Code and data to reproduce the analysis and results of this work are available at https://github.com/KULL-Centre/AF2chi. The implementation of AF2χ in localColabFold can be found at https://github.com/matteo-cagiada/AF2chi_localcolabfold

## Acknowledgments

We are grateful to Yann Vander Meersche and Tatiana Galochkina for providing the data on the computational performance of the MDatlas simulations used in our analyses. The research was supported by the PRISM (Protein Interactions and Stability in Medicine and Genomics) centre funded by the Novo Nordisk Foundation (NNF18OC0033950, to K.L.-L.), a Novo Nordisk Foundation Postdoctoral Fellowship (NNF23OC0082912; to MC). We acknowledge access to computational resources via a grant from the Carlsberg Foundation (CF21-0392; to K.L.-L.).

## Supplementary Material

### Supplementary Tables

**Table S1.** Accuracy of AF2χ and other methods for recovering HSP ensemble χ_1_ distributions.

| $\chi_1$ - JS divergence (mean ± $\sigma$) - ALL HSP residues | | | | | | |
| --- | --- | --- | --- | --- | --- | --- |
| Protein | AF2 structure | Top8000 | CHARMM36m | NMR-restrained ensemble | AF2χ prior | AF2χ |
| UBQ | 0.58 ± 0.14 | 0.47 ± 0.10 | 0.37 ± 0.10 | 0.36 ± 0.09 | **0.34 ± 0.08** | **0.34 ± 0.08** |
| HEWL | 0.51 ± 0.22 | 0.58 ± 0.08 | 0.48 ± 0.15 | | **0.41 ± 0.10** | 0.42 ± 0.10 |
| BPTI | 0.57 ± 0.17 | 0.55 ± 0.10 | 0.41 ± 0.12 | | 0.40 ± 0.10 | **0.39 ± 0.10** |

| $\chi_1$ - JS divergence (mean ± $\sigma$) - Single-well HSP residues | | | | | | |
| --- | --- | --- | --- | --- | --- | --- |
| Protein | AF2 structure | Top8000 | CHARMM36m | NMR-restrained ensemble | AF2χ prior | AF2χ |
| UBQ | 0.59 ± 0.15 | 0.54 ± 0.07 | 0.35 ± 0.11 | **0.34 ± 0.08** | **0.34 ± 0.08** | **0.34 ± 0.08** |
| HEWL | 0.47 ± 0.24 | 0.61 ± 0.07 | 0.47 ± 0.14 | | **0.43 ± 0.09** | **0.43 ± 0.09** |
| BPTI | 0.54 ± 0.19 | 0.60 ± 0.08 | 0.39 ± 0.13 | | **0.37 ± 0.09** | **0.37 ± 0.08** |

| $\chi_1$ - JS divergence (mean ± $\sigma$) - Multi-well HSP residues | | | | | | |
| --- | --- | --- | --- | --- | --- | --- |
| Protein | AF2 structure | Top8000 | CHARMM36m | NMR-restrained ensemble | AF2χ prior | AF2χ |
| UBQ | 0.59 ± 0.13 | 0.43 ± 0.10 | 0.37 ± 0.09 | 0.37 ± 0.09 | 0.35 ± 0.09 | **0.34 ± 0.07** |
| HEWL | 0.56 ± 0.18 | 0.54 ± 0.09 | 0.49 ± 0.14 | | 0.40 ± 0.11 | **0.39 ± 0.11** |
| BPTI | 0.61 ± 0.10 | 0.49 ± 0.08 | 0.45 ± 0.10 | | 0.44 ± 0.12 | **0.43 ± 0.11** |

**Table S2.** Accuracy of AF2χ and other methods for recovering HSP ensemble χ_2_ distributions.

| $\chi_2$ - JS divergence ( mean ± $\sigma$) - ALL HSP residues | | | | | | |
| --- | --- | --- | --- | --- | --- | --- |
| Protein | AF2 structure | Top8000 | CHARMM36m | NMR-restrained ensemble | AF2χ prior | AF2χ |
| UBQ | 0.61 ± 0.12 | 0.45 ± 0.10 | 0.44 ± 0.09 | 0.41 ± 0.13 | 0.38 ± 0.12 | **0.38 ± 0.09** |
| HEWL | 0.63 ± 0.18 | 0.57 ± 0.10 | 0.53 ± 0.12 | | 0.51 ± 0.12 | **0.50 ± 0.13** |
| BPTI | 0.65 ± 0.13 | 0.53 ± 0.11 | **0.49 ± 0.13** | | 0.52 ± 0.14 | 0.51 ± 0.13 |

**Table S3.** Accuracy of AF2χ and other methods for matching experimental NMR ^3^*J*-couplings.

| Protein | Dihedral | [HTML]EFEFEFAF2χ | AF2χ prior | AF2 structure | HSP ensemble | NMR-restrained ensemble | CHARMM36m | ff99sb | ff99sb-ILDN |
| --- | --- | --- | --- | --- | --- | --- | --- | --- | --- |
| BPTI | Ha-Ca-Cb-Hb | [HTML]EFEFEF1.42 ± 0.13 | 1.46 ± 0.13 | **1.16 ± 0.13** | **1.16 ± 0.10** | | 1.80 ± 0.23 | 1.78 ± 0.29 | 1.45 ± 0.13 |
| GB3 | Ha-Ca-Cb-Hb | [HTML]EFEFEF1.69 ± 0.24 | 2.06 ± 0.27 | 2.01 ± 0.37 | | | 1.71 ± 0.21 | 2.41 ± 0.45 | **1.36 ± 0.13** |
| GB3 | C-Ca-Cb-Cg | [HTML]EFEFEF0.67 ± 0.17 | 0.71 ± 0.17 | 0.89 ± 0.18 | | | 0.62 ± 0.14 | **0.59 ± 0.08** | 0.61 ± 0.13 |
| GB3 | N-Ca-Cb-Cg | [HTML]EFEFEF**0.20 ± 0.02** | 0.22 ± 0.03 | 0.29 ± 0.04 | | | **0.20 ± 0.02** | 0.28 ± 0.04 | 0.26 ± 0.04 |
| UBQ | Ha-Ca-Cb-Hb | [HTML]EFEFEF1.71 ± 0.18 | 2.02 ± 0.20 | 2.30 ± 0.22 | **1.39 ± 0.15** | 1.90 ± 0.15 | 1.65 ± 0.16 | 2.54 ± 0.24 | 1.92 ± 0.17 |
| UBQ | C-Ca-Cb-Cg | [HTML]EFEFEF0.82 ± 0.11 | 0.87 ± 0.11 | 1.00 ± 0.11 | **0.71 ± 0.14** | 0.85 ± 0.12 | 0.83 ± 0.09 | 1.07 ± 0.22 | 0.82 ± 0.13 |
| UBQ | N-Ca-Cb-Cg | [HTML]EFEFEF0.25 ± 0.04 | 0.29 ± 0.05 | 0.31 ± 0.05 | 0.23 ± 0.04 | **0.21 ± 0.03** | 0.31 ± 0.06 | 0.46 ± 0.09 | 0.31 ± 0.06 |
| HEWL | C-Ca-Cb-Hb | [HTML]EFEFEF2.02 ± 0.40 | 2.00 ± 0.40 | **1.95 ± 0.44** | 1.99 ± 0.43 | | 2.38 ± 0.39 | 2.71 ± 0.37 | 2.69 ± 0.36 |
| HEWL | Ha-Ca-Cb-Hb | [HTML]EFEFEF2.48 ± 0.14 | 2.51 ± 0.13 | **2.13 ± 0.12** | 2.20 ± 0.13 | | 3.31 ± 0.27 | 3.59 ± 0.32 | 2.92 ± 0.17 |

**Table S4.** Accuracy of AF2χ and other methods for matching experimental NMR methyl 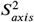 order parameters.

| Protein | **AF2χ** | AF2χ prior | CHARMM36m | Null model |
| --- | --- | --- | --- | --- |
| A_LBP | 0.64 ± 0.11 | 0.47 ± 0.14 | **0.71 ± 0.10** | 0.66 ± 0.09 |
| FNfn10 | 0.65 ± 0.06 | 0.44 ± 0.12 | **0.69 ± 0.09** | 0.47 ± 0.10 |
| M_FABP | **0.62 ± 0.09** | 0.52 ± 0.11 | **0.62 ± 0.10** | 0.44 ± 0.13 |
| A3D | 0.75 ± 0.07 | 0.75 ± 0.07 | 0.47 ± 0.14 | **0.80 ± 0.06** |
| Fyn_SH3 | 0.53 ± 0.15 | 0.38 ± 0.19 | **0.82 ± 0.08** | 0.69 ± 0.13 |
| TNfn3 | 0.55 ± 0.11 | 0.56 ± 0.10 | 0.49 ± 0.12 | **0.60 ± 0.10** |

**Table S5.** Accuracy of AF2χ and generative structural ensemble methods for recovering HSP ensemble χ_1_ distributions.

| **$\chi_1$ - JS divergences ( mean ± $\sigma$) - ALL HSP residues** |  |  |  |  |  |  |  |
| --- | --- | --- | --- | --- | --- | --- | --- |
| | AF2 structure | AF2 dropout | Seqdance | MDgen | BioEmu + Hpacker | aSAMt | AF2χ |
| UBQ | 0.58 ± 0.14 | 0.48 ± 0.13 | 0.73 ± 0.07 | 0.40 ± 0.17 | 0.39 ± 0.09 | 0.39 ± 0.09 | **0.34 ± 0.08** |
| HEWL | 0.51 ± 0.22 | 0.46 ± 0.20 | 0.74 ± 0.04 | 0.47 ± 0.18 | 0.48 ± 0.12 | 0.49 ± 0.12 | **0.42 ± 0.10** |
| BPTI | 0.58 ± 0.17 | 0.44 ± 0.14 | 0.74 ± 0.07 | 0.40 ± 0.15 | 0.42 ± 0.12 | 0.49 ± 0.12 | **0.39 ± 0.10** |

| **$\chi_1$ - JS divergences ( mean ± $\sigma$) - Single well HSP residues** |  |  |  |  |  |  |  |
| --- | --- | --- | --- | --- | --- | --- | --- |
| | AF2 structure | AF2 dropout | Seqdance | MDgen | BioEmu + Hpacker | aSAMt | AF2χ |
| UBQ | 0.55 ± 0.15 | 0.41 ± 0.11 | 0.78 ± 0.05 | 0.36 ± 0.17 | 0.36 ± 0.12 | 0.39 ± 0.09 | **0.34 ± 0.08** |
| HEWL | 0.47 ± 0.24 | 0.41 ± 0.21 | 0.77 ± 0.02 | 0.45 ± 0.18 | 0.45 ± 0.12 | 0.49 ± 0.12 | **0.43 ± 0.09** |
| BPTI | 0.55 ± 0.19 | 0.38 ± 0.13 | 0.76 ± 0.05 | **0.37 ± 0.15** | 0.40 ± 0.13 | 0.49 ± 0.09 | **0.37 ± 0.08** |

| **$\chi_1$ - JS divergences ( mean ± $\sigma$) - Multi wells HSP residues** |  |  |  |  |  |  |  |
| --- | --- | --- | --- | --- | --- | --- | --- |
| | AF2 structure | AF2 dropout | Seqdance | MDgen | BioEmu + Hpacker | aSAMt | AF2χ |
| UBQ | 0.59 ± 0.13 | 0.53 ± 0.11 | 0.71 ± 0.07 | 0.42 ± 0.07 | 0.40 ± 0.10 | 0.39 ± 0.09 | **0.34 ± 0.07** |
| HEWL | 0.56 ± 0.18 | 0.51 ± 0.18 | 0.70 ± 0.05 | 0.50 ± 0.17 | 0.50 ± 0.11 | 0.48 ± 0.12 | **0.39 ± 0.11** |
| BPTI | 0.63 ± 0.10 | 0.53 ± 0.11 | 0.69 ± 0.07 | 0.45 ± 0.15 | 0.46 ± 0.10 | 0.48 ± 0.09 | **0.43 ± 0.11** |

**Table S6.** Accuracy of AF2χ and generative structural ensemble methods for matching experimental NMR ^3^*J*-coupling.

| Protein | Dihedral | **AF2χ** | MDgen | Bioemu + Hpacker | aSAMt |
| --- | --- | --- | --- | --- | --- |
| BPTI | Ha-Ca-Cb-Hb | **1.42 ± 0.13** | 1.73 ± 0.33 | 1.80 ± 0.15 | 2.51 ± 0.24 |
| UBQ | Ha-Ca-Cb-Hb | 1.71 ± 0.18 | 2.31 ± 0.21 | **1.60 ± 0.16** | 1.94 ± 0.14 |
| UBQ | C-Ca-Cb-Cg | 0.82 ± 0.11 | 1.312 ± 0.13 | **0.65 ± 0.10** | 0.81 ± 0.10 |
| UBQ | N-Ca-Cb-Cg | **0.25 ± 0.04** | 0.47 ± 0.15 | 0.28 ± 0.04 | 0.32 ± 0.05 |
| HEWL | C-Ca-Cb-Hb | **2.02 ± 0.40** | 2.30 ± 0.13 | 2.2 ± 0.40 | 2.25 ± 0.40 |
| HEWL | Ha-Ca-Cb-Hb | **2.48 ± 0.14** | 2.82 ± 0.33 | 2.91 ± 0.21 | 3.08 ± 0.28 |

### Supplementary Figures

**Figure S1.**
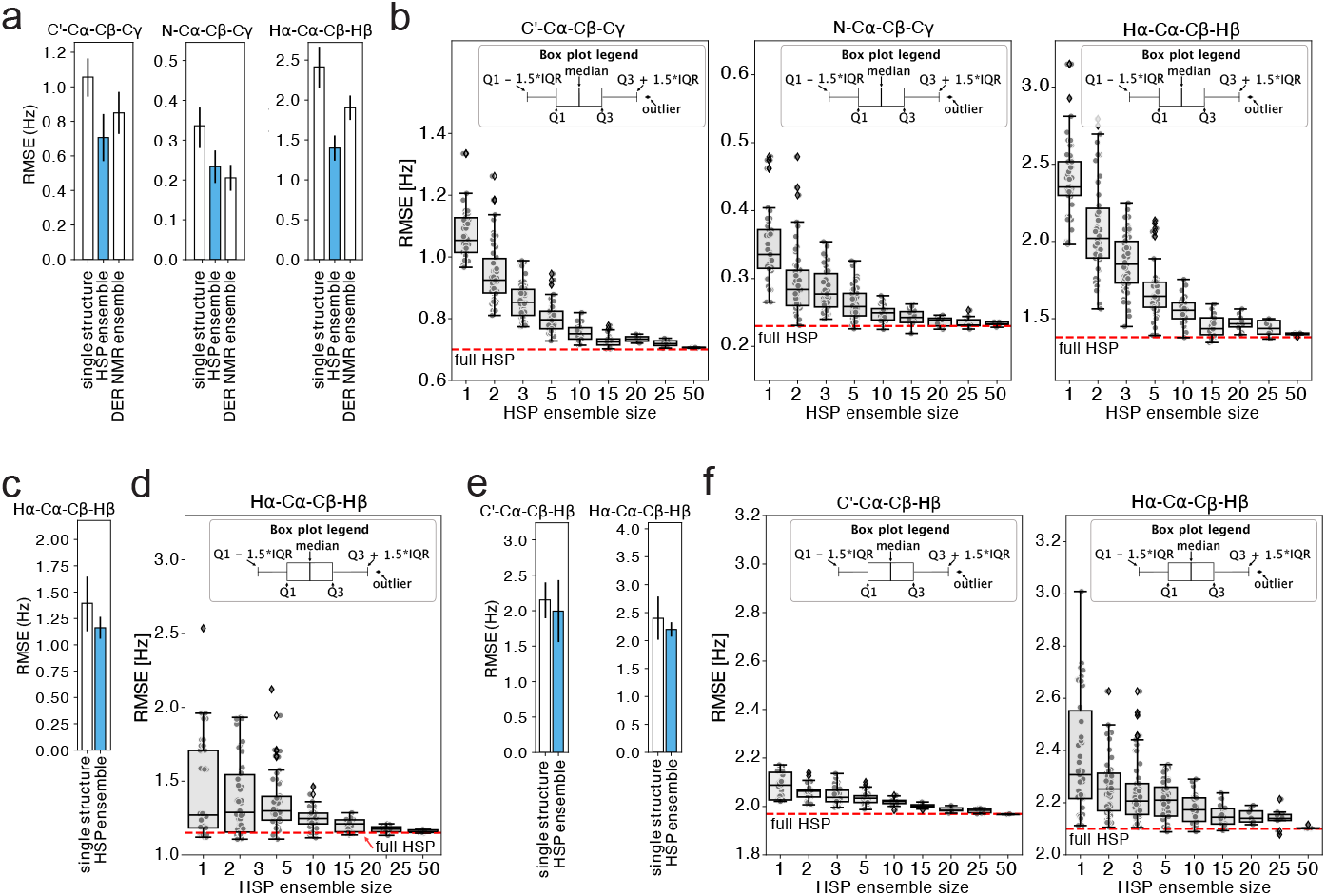
HSP ensemble accuracy for ^3^*J*-couplings and effect of HSP ensemble size. **(a**,**c**,**e)** Root mean square error (RMSE) to experimental ^3^*J*-couplings for ^3^*J*-couplings calculated from the HSP ensemble (light blue bar) and, for comparison, for ^3^*J*-couplings calculated from the single structure (AF2 prediction) and the DER ensemble (white bar). The black vertical line at the top of each column shows the sample standard deviation of bootstrapped distributions. **(b**,**d**,**f)** RMSEs (y-axis) as a function of the number of structures in the HSP ensemble (x-axis) used to calculate the different UBQ ^3^*J*-couplings. The box plots show the RMSE statistics as a result of bootstrapping structures from the full HSP ensemble to different ensemble sizes. The red horizontal line in each plot represents the accuracy of the full HSP ensemble.

**Figure S2.**
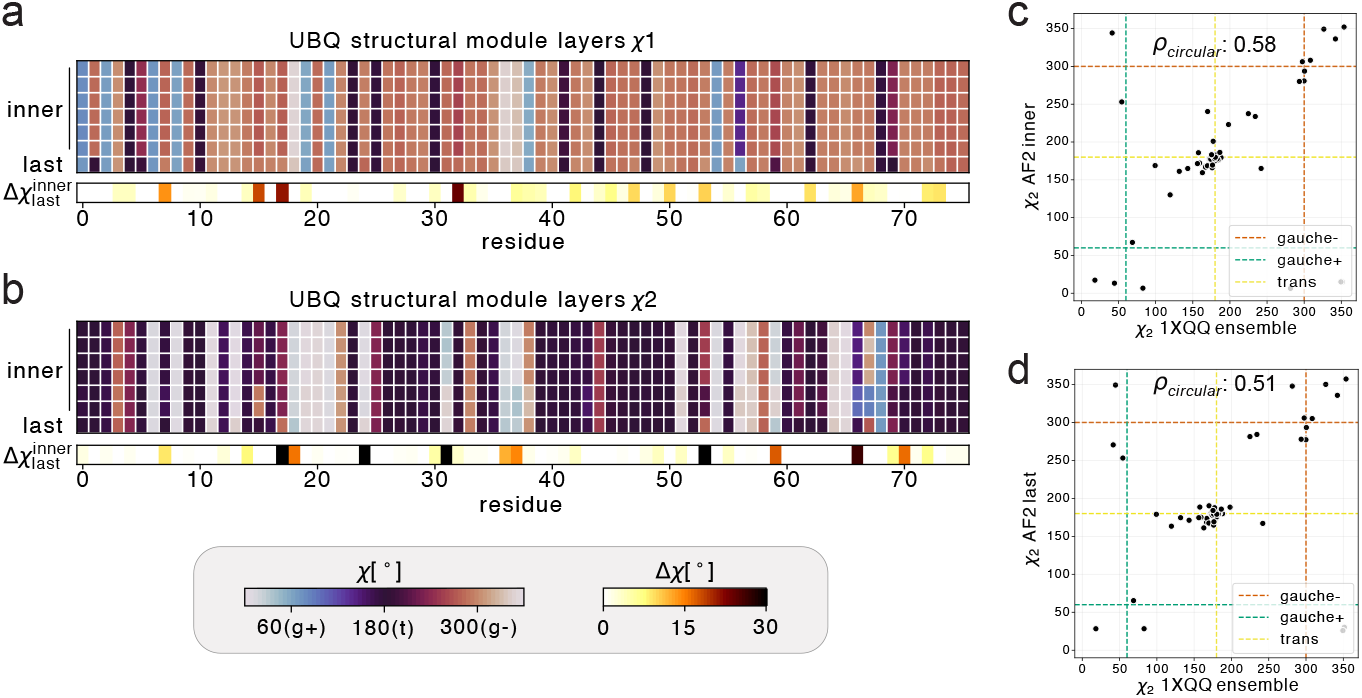
Differences in χ-angle predictions in the AF2 structure module and comparison between AF2 and DER ensemble χ_1_ predictions. Inner- and last-layer AF2 predictions of **(a)** χ_1_ and **(b)** χ_2_ for all residues in UBQ. The top heatmap shows the χ-angle prediction in a given layer (y-axis) for each residue (x-axis). The bottom heatmap shows the difference, Δχ, between the averaged inner-layer χ-angles and the last-layer χ-angle for **(a)** χ_1_ and **(b)** χ_2_ for all residues in UBQ. **(c**,**d**) Correlation between the averaged χ_2_-angles from the DER ensemble (PDB: 1XQQ) and the **(c)** averaged χ_2_ values from AF2 structure module inner layers and **(d)** χ_2_ values from the structure module last layer.

**Figure S3.**
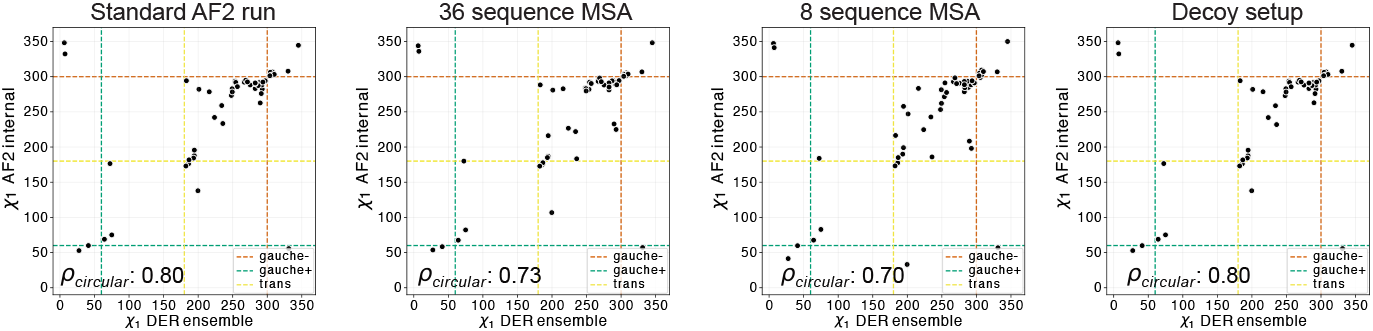
AF2 parameter test. Benchmark of different AF2 setups to generate an optimal target ensemble average for BME reweighting in AF2χ. Each plot shows the correlation of averaged inner-layer χ_1_-angles from the AF2 structural module (y-axis) and averaged χ_1_-angles from the DER ensemble (x-axis) for all the DER ensemble multi-well residues. The four different setups tested are: AF2 standard parameter run (left), AF2 with MSA subsample (32 sequences middle left, 8 sequences middle right) and the decoy setup (right). The main χ_1_-angle states are shown as dotted lines and the circular Pearson’s correlation coefficient is shown in the bottom left corner.

**Figure S4.**
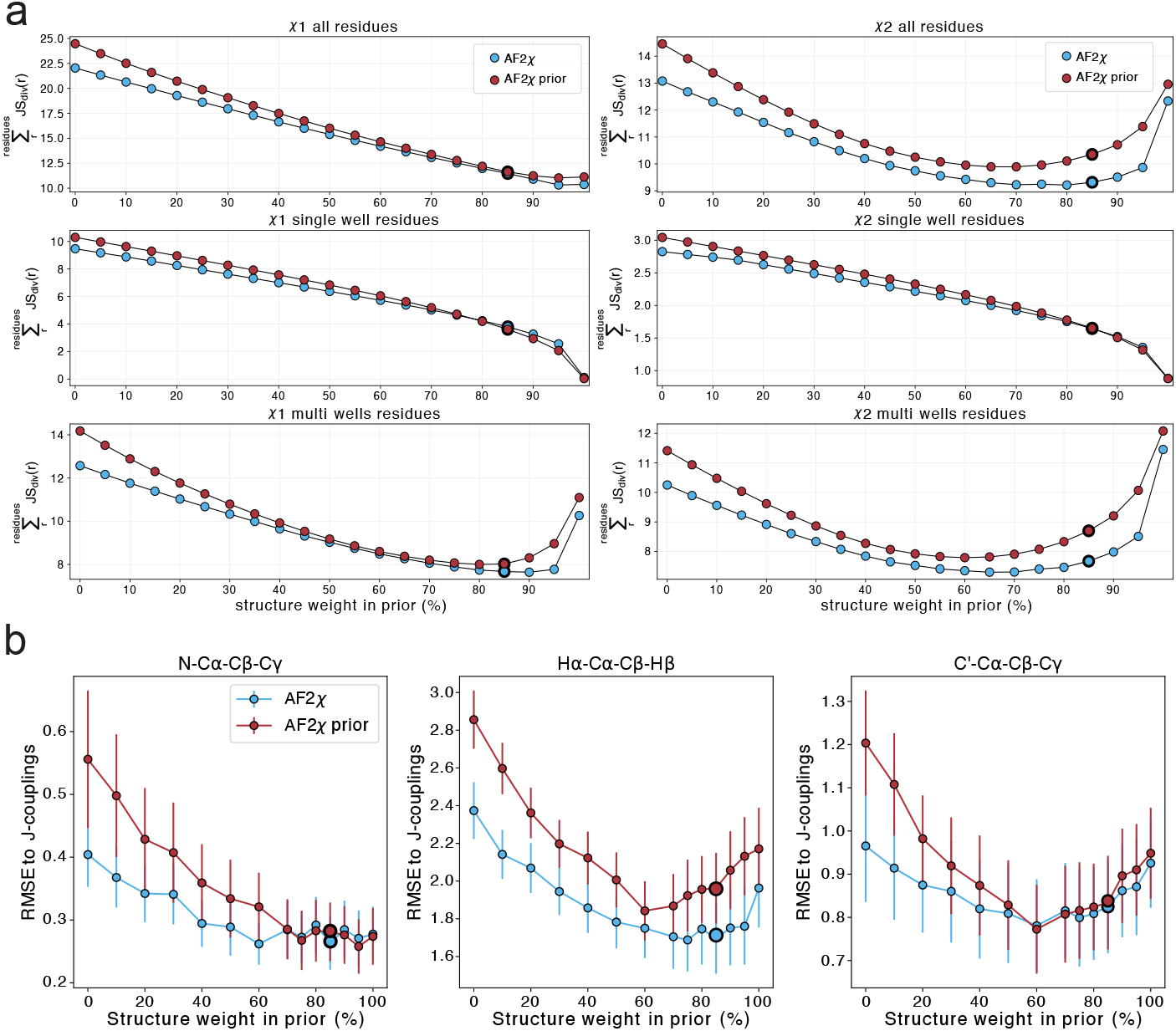
Selection of the optimal mixing ratio between AF2 structure χ-angles and Top8000 distributions to create prior distributions for AF2χ. **(a)** Sum of the JS divergences (y-axis) between AF2χ and HSP χ-angle distributions for UBQ. Top plots show the JS-divergence values for χ_1_ (left) and χ_2_ (right) for all residues, while the middle and bottom plots show the correlation for the subsets of single- (middle) and multi-well (bottom) residues based on the HSP ensemble. **(b)** RMSE between ^3^*J*-couplings calculated from AF2χ structural ensembles and experimental ^3^*J*-couplings that report on χ_1_-angles for UBQ. The vertical line on each marker shows the sample standard deviation of bootstrapped distributions. **(a**,**b)** The mixing ratio is shown on the x-axis (corresponding to *α* in equation 1), with a prior consisting only of Top8000 distributions on the extreme left or a Gaussian distribution around AF2 last-layer values on the extreme right. Red denotes prior distributions and blue denotes posterior distributions output by AF2χ. The optimal mixing ratio (85% AF2 last-layer distribution, 15% Top8000 distribution) is shown as markers with thicker black edges.

**Figure S5.**
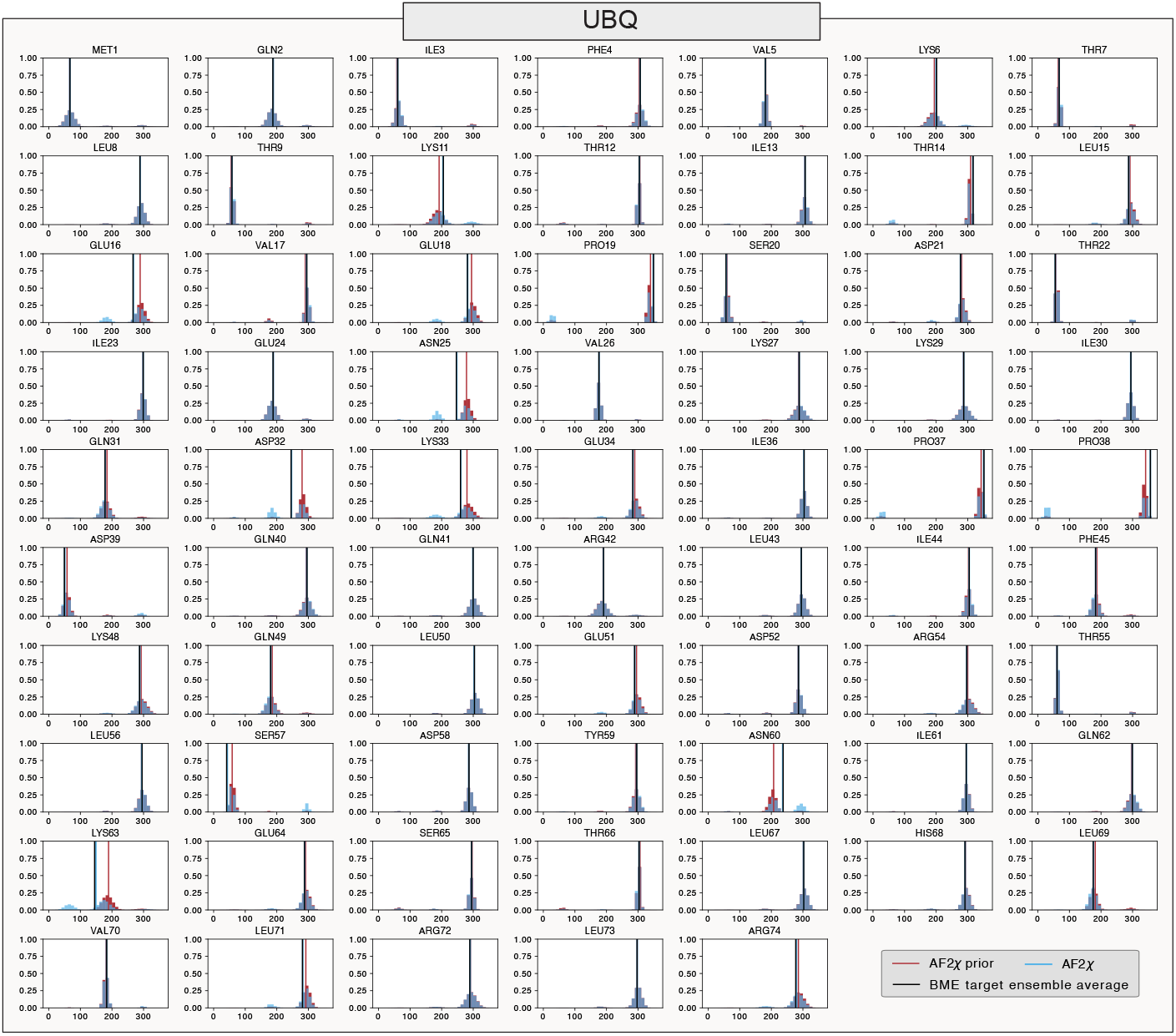
Reweighting results for UBQ χ_1_ rotamer distributions. Comparison of χ_1_ rotamer distributions from the AF2χ prior (red) and AF2χ final output (light blue). Black lines report the average inner-layer χ_1_ from the AF2 structure module used as the target average for BME reweighting. Red lines show the average distribution value (circular mean) for the AF2χ BME prior, and blue lines the circular averaged values for the AF2χ reweighted distribution.

**Figure S6.**
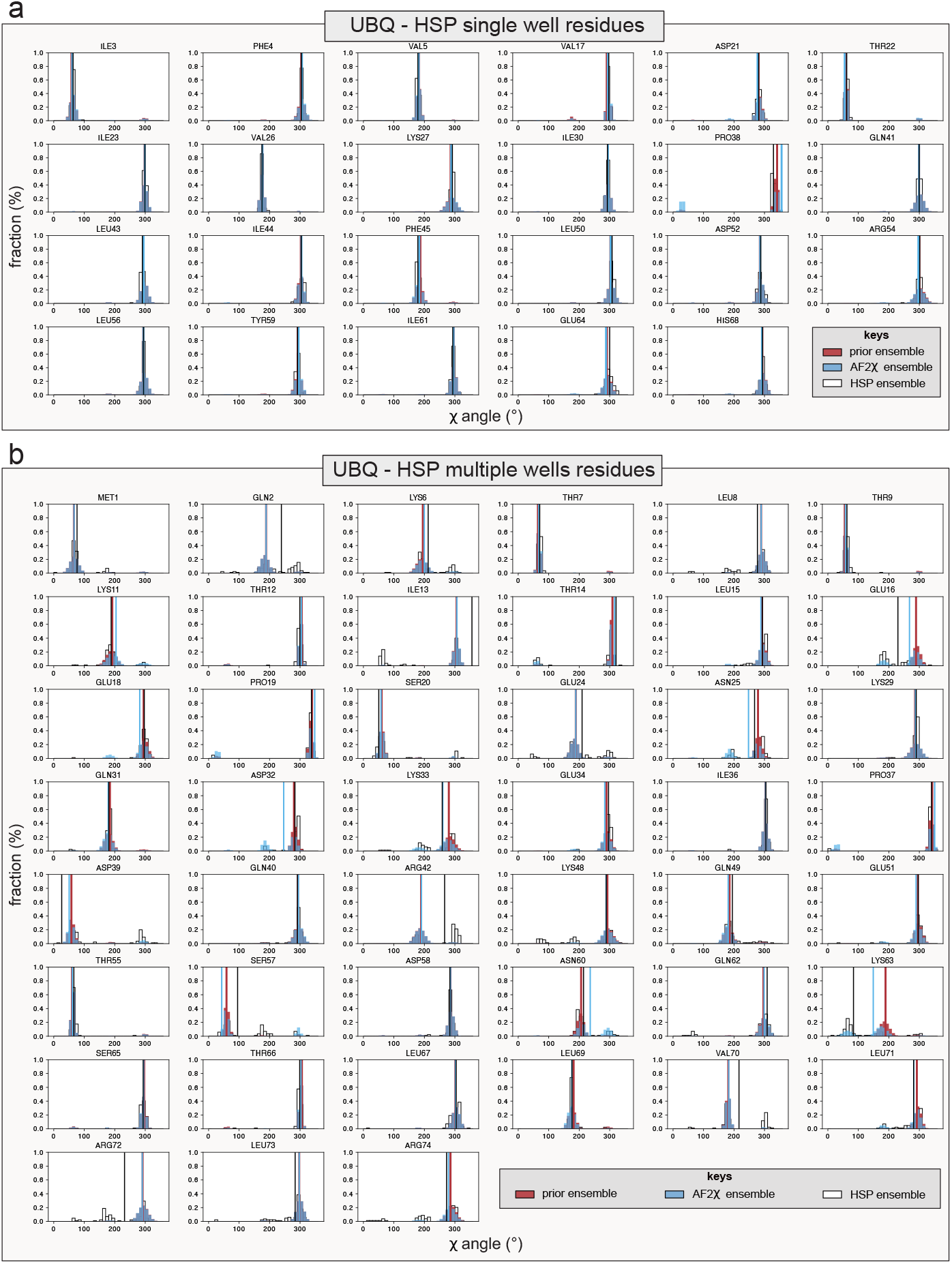
UBQ χ_1_ rotamer distributions. **(a**,**b)** Comparison of UBQ χ_1_-angle distributions for **(a)** single- and **(b)** multi-well residues based on the HSP ensemble. AF2χ prior distributions are shown in red, final AF2χ distributions are shown in blue and HSP ensemble distributions are shown with a white-black contour. Circular means are reported as vertical lines in the same colour as the corresponding distribution.

**Figure S7.**
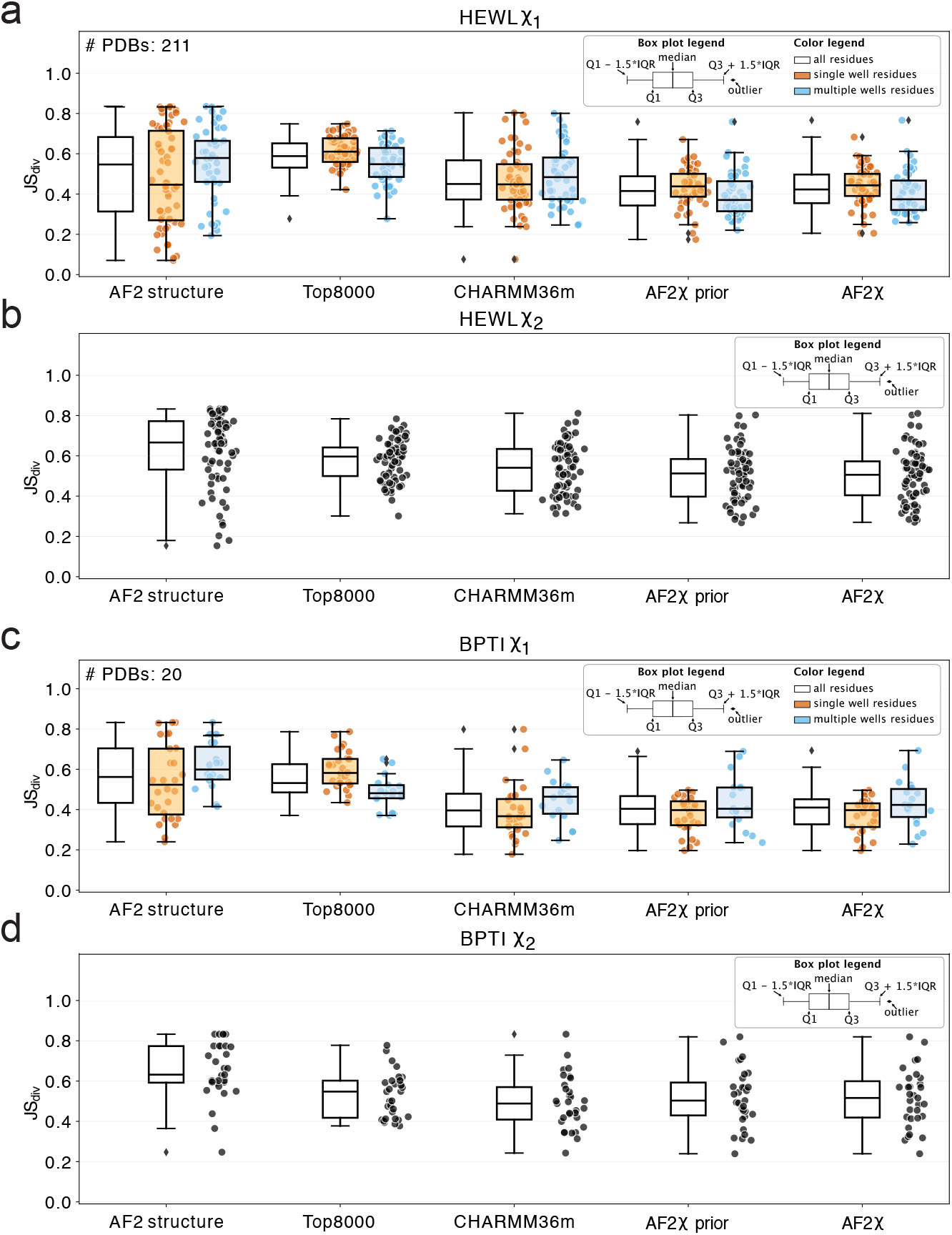
Validation of HEWL and BPTI AF2χ rotamer distributions using HSP ensembles. **(a**,**c)** JS divergences between AF2χ and HSP ensemble χ_1_-angle distributions for **(a)** HEWL and **(c)** BPTI. The left box plot (white) shows the JS divergence statistics for all residues in the protein, the middle box plot (orange) for single-well residues based on the HSP ensemble, and the right box plot (light blue) for multi-well based on the HSP ensemble. The individual JS-divergence values are shown as dots below the single- and multi-well box plots for each tested method. **(b**,**d)** JS divergences between AF2χ and HSP ensemble χ_2_-angle distributions for **(b)** HEWL and **(d)** BPTI., with a box plot showing the JS divergence statistics on the right and the individual JS-divergence values on the left for each tested method.

**Figure S8.**
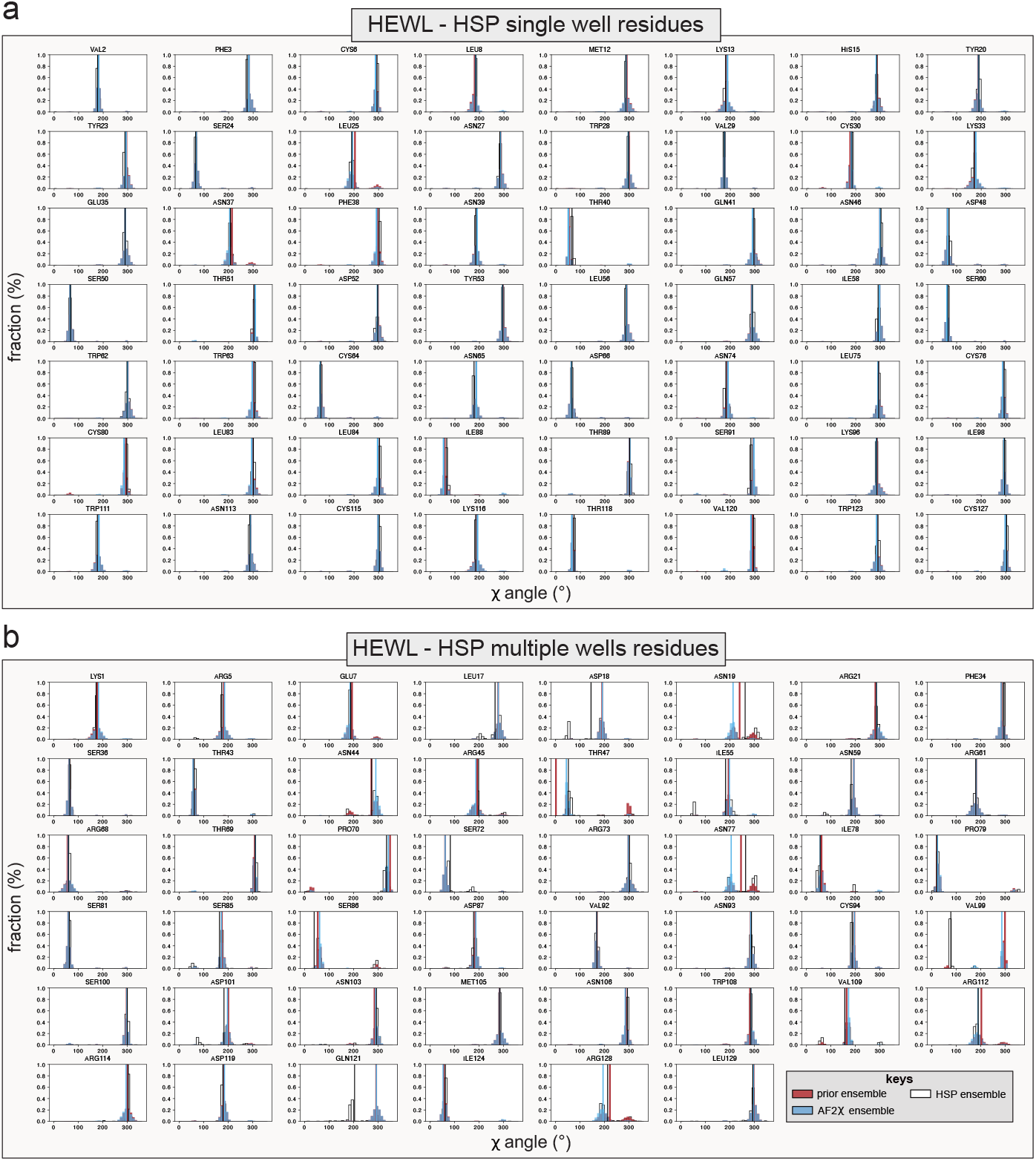
HEWL χ_1_ rotamer distributions. **(a**,**b)** Comparison of HEWL χ_1_-angle distributions for **(a)** single- and **(b)** multi-well residues based on the HSP ensemble. AF2χ prior distributions are shown in red, final AF2χ distributions are shown in blue and HSP ensemble distributions are shown with a white-black contour. Circular means are reported as vertical lines in the same colour as the corresponding distribution.

**Figure S9.**
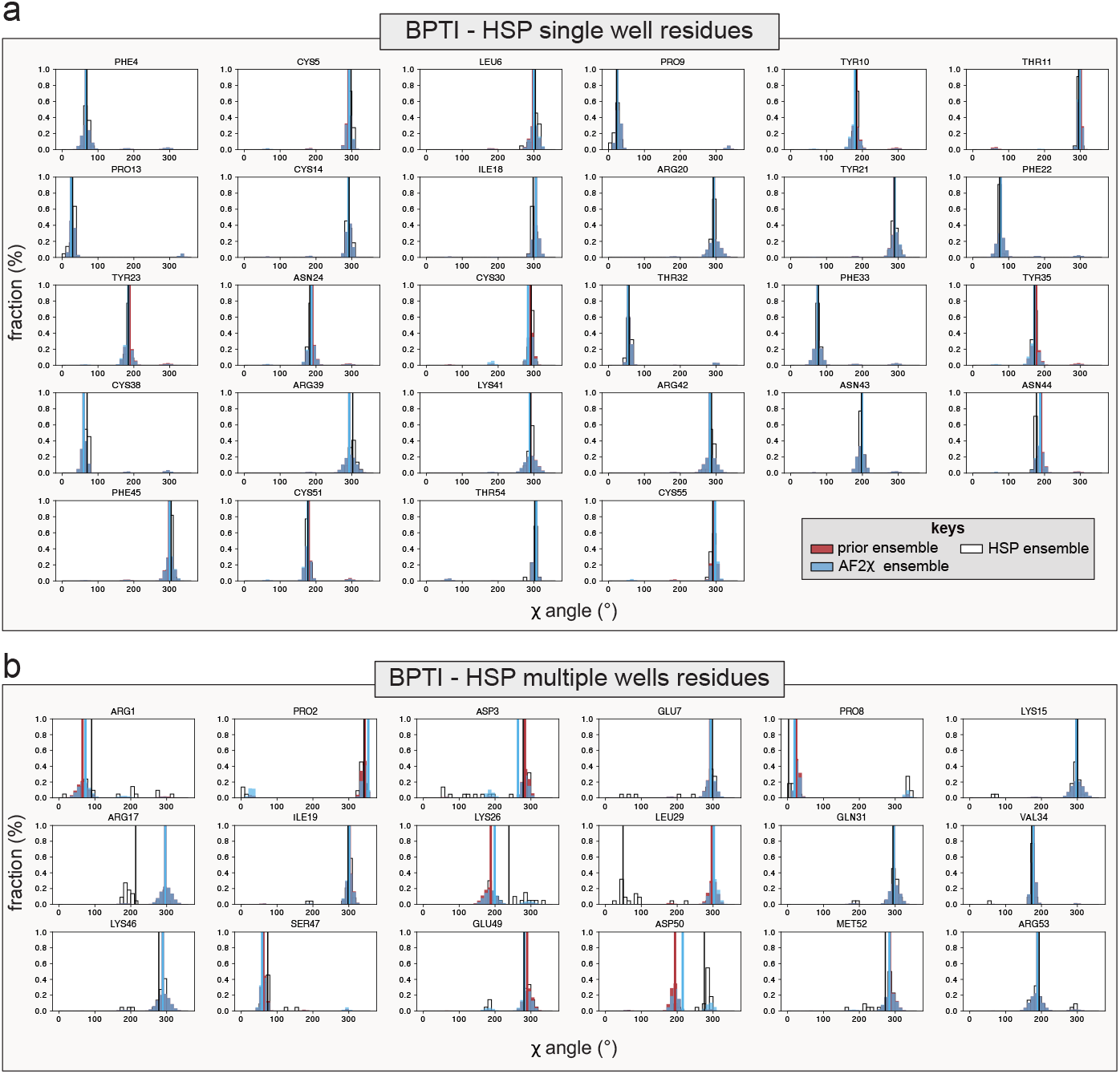
BPTI χ_1_ rotamer distributions. **(a**,**b)** Comparison of BPTI χ_1_-angle distributions for **(a)** single- and **(b)** multi-well residues based on the HSP ensemble. AF2χ prior distributions are shown in red, final AF2χ distributions are shown in blue and HSP ensemble distributions are shown with a white-black contour. Circular means are reported as vertical lines in the same colour as the corresponding distribution.

**Figure S10.**
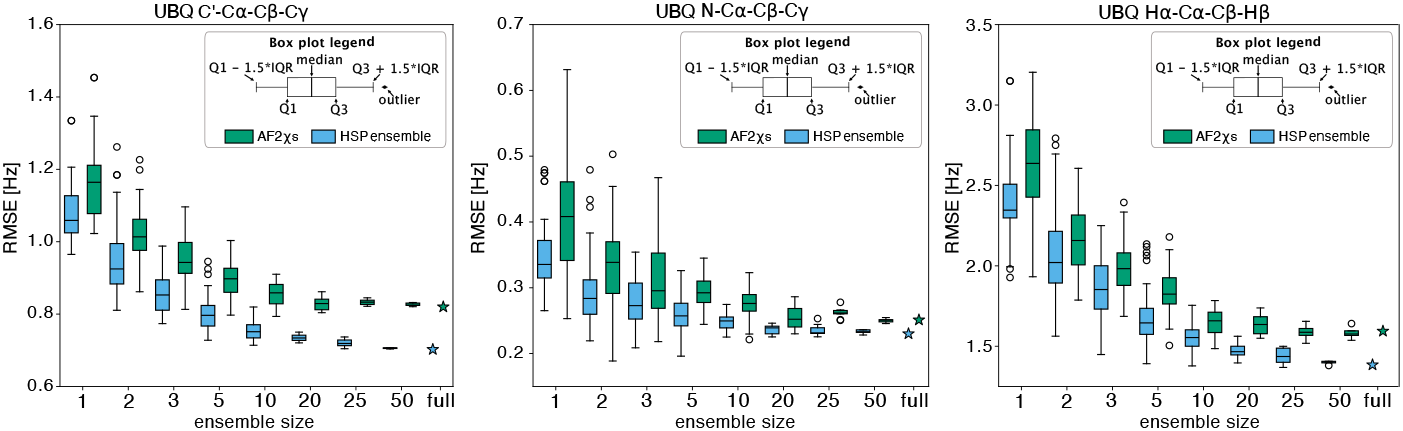
Effect of AF2χ ensemble size on the prediction of ^3^*J*-couplings. Root mean square error (RMSE, y-axis) to the experimental UBQ ^3^*J*-couplings for ^3^*J*-couplings calculated from AF2χ (blue) and HSP (green) ensembles as a function of the number of structures in the ensemble (x-axis). The box plots show the RMSE statistics as a result of bootstrapping the ensemble structures and the star marker represents the RMSE given by the full ensemble.

**Figure S11.**
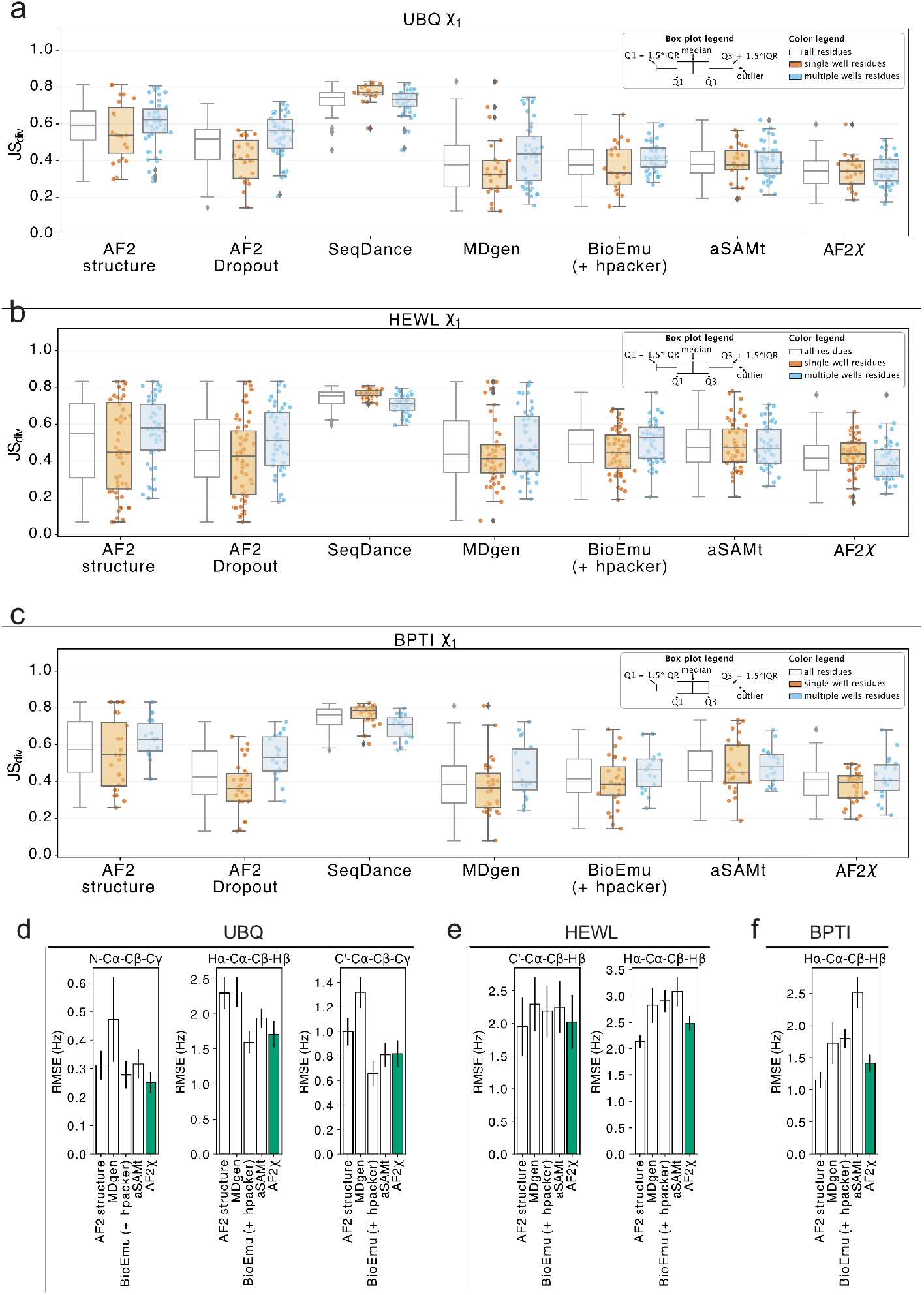
Comparison of AF2χ accuracy with other methods to predict conformational ensembles. **(a**,**b**,**c)** Comparison, using the JS divergence (y-axis) to HSP ensemble χ_1_-angle distributions, between AF2χ rotamer distributions and those from recently developed methods for structural ensembles of proteins. Results are reported for **(a)** UBQ, **(b)** HEWL, and **(c)** BPTI. The left box plot (white) shows the JS divergence statistics for all residues in the protein, the middle box plot (orange) for the single-well residues and the right box plot (blue) for the multi-well residues based on the HSP ensemble. The individual JS-divergence values are shown as dots below the single- and multi-well plots. **(d**,**e**,**f)** RMSE (y-axis) to experimental ^3^*J*-couplings on χ_1_ for the protocols tested (x-axis). AF2χ RMSEs are shown as green bars. The black vertical line at the top of each column shows the sample standard deviation of bootstrapped distributions.

**Figure S12.**
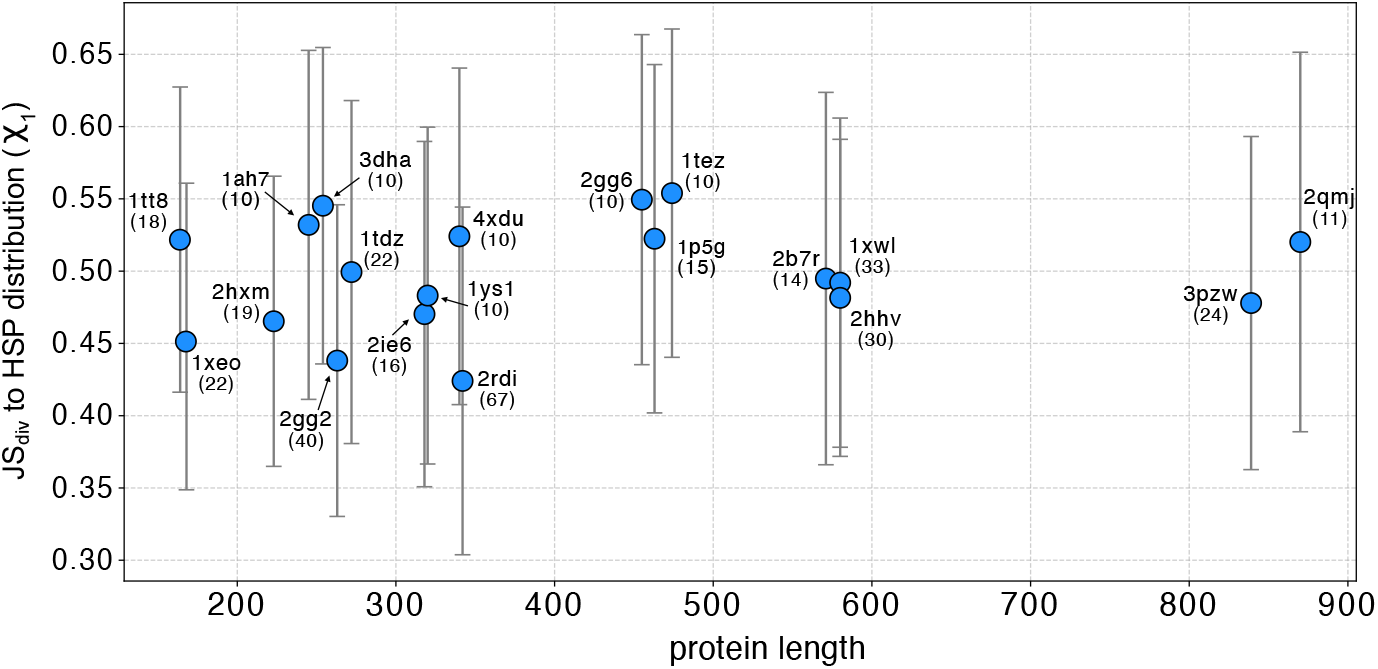
Assessment of accuracy of AF2χ for proteins with different sizes. The figure shows the protein-averaged JS divergence (y-axis) to HSP χ_1_-angle distributions for AF2χ output distributions, with protein length reported on the x-axis. The error bars represent the standard deviation for each protein, which were selected as the MDatlas proteins for which we could generate HSP ensembles. The PDB used for each inference and the number of structures in the HSP ensemble are shown next to the average (blue point).

**Figure S13.**
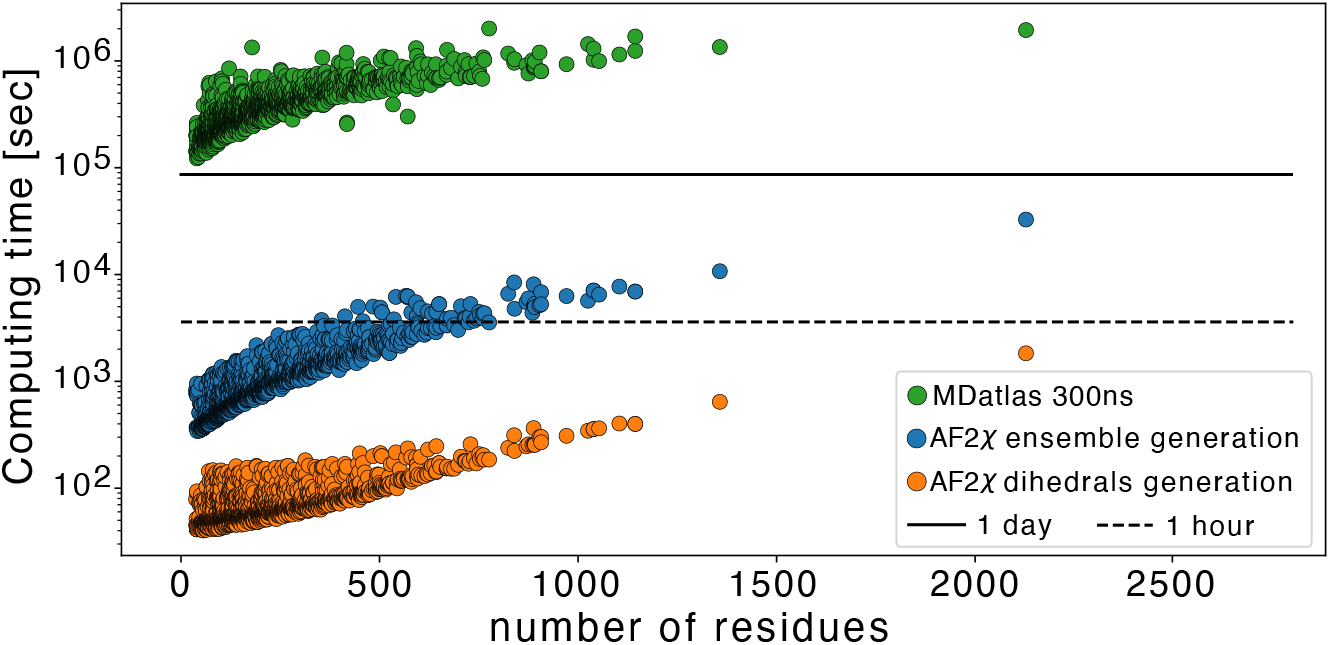
Computational time for AF2χ and MD simulations. Computational times (y-axis, seconds) of AF2χ in generating dihedral distributions (orange dots) and structural ensembles (blue dots) for all proteins in the ATLAS MD dataset as a function of their size (x-axis). For comparison, we also show the computational times used for running the ATLAS MD simulations (green dots). The black dotted and solid lines show the location for one hour and one day respectively on the y-axis.

**Figure S14.**
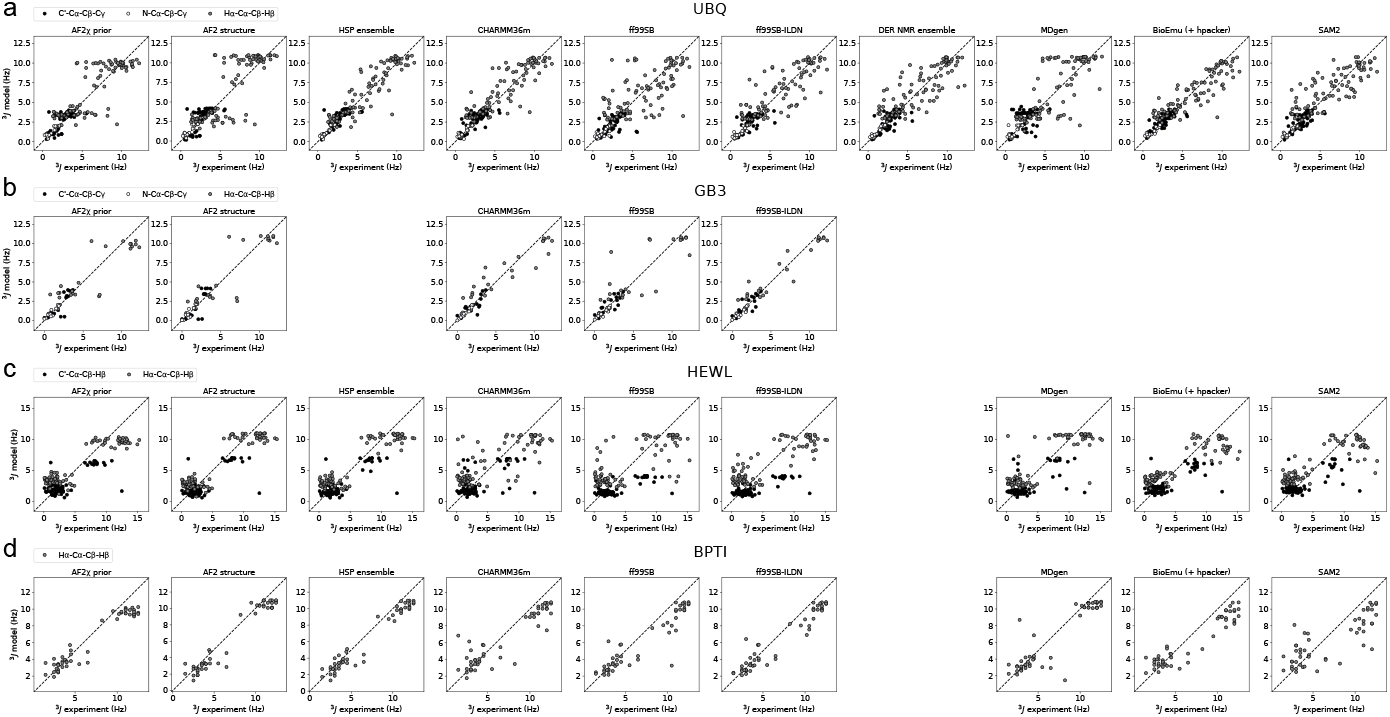
Benchmarking using ^3^*J*-couplings. Comparison of experimental NMR ^3^*J*-couplings with ^3^*J*-couplings calculated from different structural models using Karplus equations for **(a)** UBQ, **(b)** GB3, **(c)** HEWL, and **(d)** BPTI. Points are coloured according to the χ_1_ dihedral probed by the ^3^*J*-couplings, and a corresponding legend is shown for each protein.

**Figure S15.**
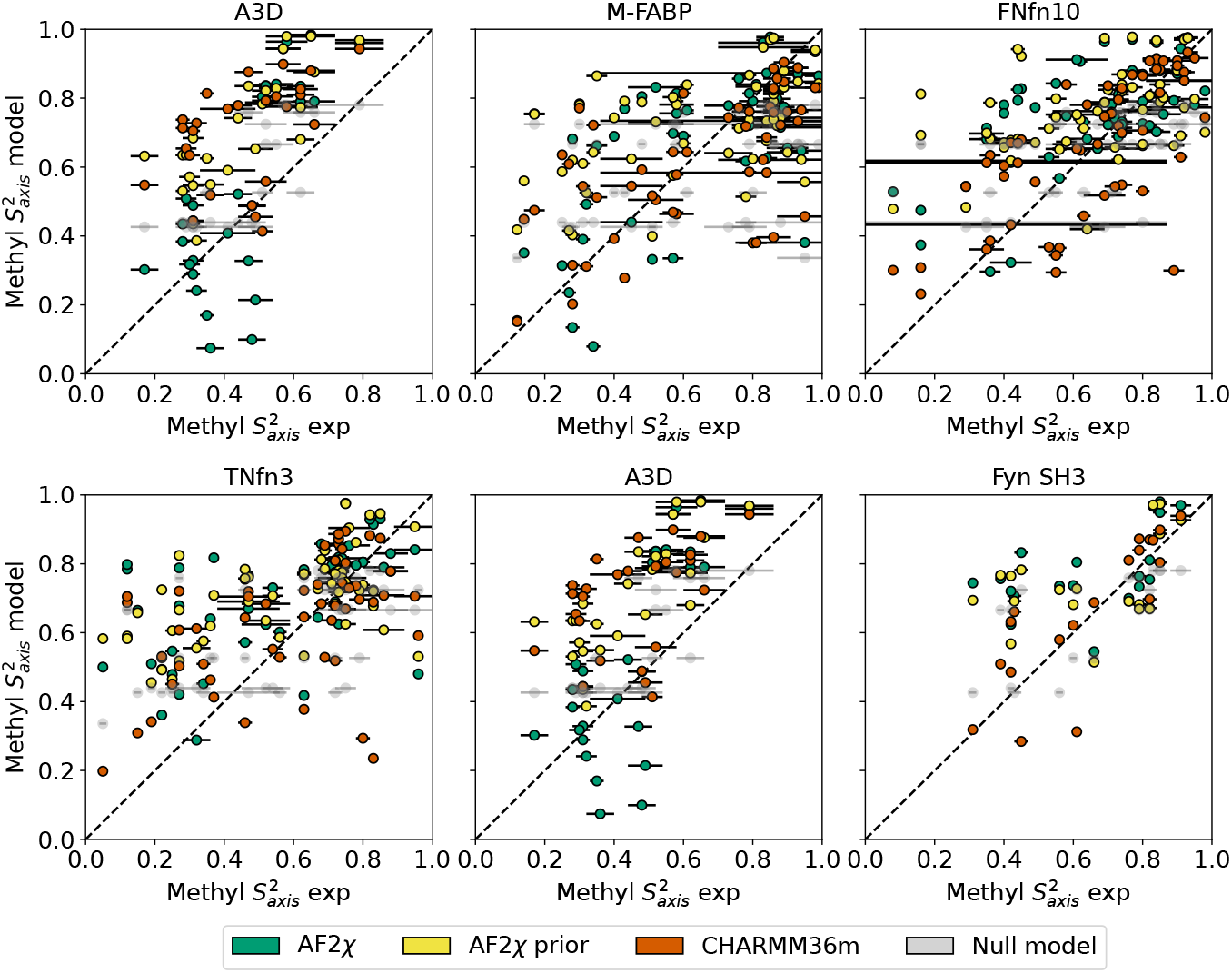
Benchmarking using methyl 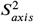 order parameters. Comparison of experimental NMR methyl 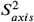 order parameters with methyl 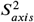 order parameters calculated from AF2χ structural ensembles (green), AF2χ prior structural ensembles (yellow), MD simulations with CHARMM36m (red), and a null model (grey) for six proteins.

